# The ontogeny of repeatability: measurement choices shape individuality estimates in a clonal fish

**DOI:** 10.64898/2026.10.15.759344

**Authors:** Nicole Walasek, Ulrike Scherer, Max Wolf, Sean M. Ehlman

## Abstract

Consistent among-individual differences in behavior across time and contexts (‘individuality’, ‘animal personality’) are increasingly recognized as playing an important role in both the ecology and evolution of species. Such individuality is not static: individual differences emerge, shift, and crystallize across ontogeny, shaped by developmental plasticity, stochastic processes, and gene-environment interactions. Despite growing awareness of this developmental dimension, the field has relied predominantly on snapshot designs with few repeated measures, raising the question of how much reported repeatability estimates reflect biology versus methodological choices. Here, we leverage high-resolution tracking data from 45 clonal Amazon mollies (*Poecilia formosa*) recorded at 5 frames per second across the first 28 days of life to systematically examine how temporal aggregation, sampling rate, frame rate, and the number and placement of measurement days affect repeatability estimates and their underlying variance components. We find that repeatability increases with higher temporal aggregation, decreases with lower sampling rates, and varies substantially depending on the developmental stage sampled. These results demonstrate that commonly used measurement practices can substantially distort repeatability estimates, and that conclusions about behavioral consistency may be as much a product of study design as of biology. We close with practical recommendations for researchers studying individuality across development.

## Introduction

Consistent individual differences in behavior are not mere noise: they influence survival, reproduction, social interactions, and population dynamics, for example through their effects on foraging, predator avoidance, and dispersal (Bro-Jørgensen et al., 2019; Réale et al., 2007). Researchers have become increasingly interested in understanding the sources of such ‘individuality’ (Sánchez-Tójar et al., 2022; Sih et al., 2004), recognizing that they can be both the fuel for and the outcome of natural selection (Dingemanse & Réale, 2005; Grunst & Grunst, 2024; Laskowski, Chang, et al., 2022; Réale et al., 2007; Scherer et al., 2023; Wolf & Weissing, 2012).

Individual differences in behavior can arise from multiple interacting sources. Genetic differences provide a heritable substrate for behavioral divergence (Mulder et al., 2016; van Oers & Mueller, 2010), while developmental plasticity—the ability of a single genotype to produce different phenotypes depending on environmental conditions experienced during ontogeny (West-Eberhard, 2003)—can generate lasting among-individual differences even when genetic backgrounds are similar (Stamps & Groothuis, 2010a; Trillmich et al., 2018). Yet, even genetically identical individuals reared under standardized conditions reliably differ in their behavior (de Bivort et al., 2022; Ehlman et al., 2025; Freund et al., 2013, 2015; Laskowski, Bierbach, et al., 2022; Scherer et al., 2023), suggesting that individuality can also emerge through developmental processes without systematic genetic or environmental differences (Bruijning et al., 2020; Frankenhuis & Panchanathan, 2011; Walasek et al., 2026). One such mechanism is developmental noise: stochastic fluctuations in gene expression and cellular processes that can generate phenotypic divergence among otherwise similar individuals (Honegger & de Bivort, 2018; Molenaar et al., 1993; Raj & van Oudenaarden, 2008; Vogt et al., 2008).

### The development of individuality

Crucially, these sources do not act once and fix behavior permanently. Early individuality research and plasticity research were often framed as opposing perspectives: the former emphasized among-individual consistency, while the latter emphasized within-individual change across time or context. We now recognize that these processes not only coexist but interact: individuals can differ in their average behavioral expression and simultaneously in how much and in what direction their behavior changes across contexts or over time. These dynamics can be captured by behavioral reaction norms, which decompose behavior into an individual intercept (personality) and slope (plasticity), with among-individual differences in slopes contributing to among-individual differences in trait expression across development (Dingemanse et al., 2010; Dingemanse & Dochtermann, 2013; Groothuis & Trillmich, 2011; Stamps & Groothuis, 2010a, 2010b; Trillmich et al., 2018). Individuality should therefore not be expected to be static across ontogeny; rather, consistent individual differences can emerge, shift, and crystallize at different rates across developmental stages. Empirically, behavioral consistency has been shown to vary across developmental stages in multiple taxa. In zebra finches, fearlessness and exploration are consistent within and across all developmental stages, while activity and aggression are only consistent at specific stages (Wuerz & Krüger, 2015); in mosquitofish, individual consistency of activity and risk-taking increases as individuals mature (Polverino et al., 2016); and in Siberian hamsters, social behavior is consistent within the juvenile stage but not across developmental stages (Adaniya et al., 2021). Yet empirical studies equipped to track the unfolding of individuality across ontogeny remain rare.

### Quantifying individuality: repeatability and its challenges

Individuality is most commonly quantified through *repeatability*: the proportion of total phenotypic variance attributable to consistent among-individual differences (Hayes & Jenkins, 1997; Nakagawa & Schielzeth, 2010; Shrout & Fleiss, 1979). Interest in repeatability has grown substantially: a systematic search on the Web of Science for empirical papers on individuality and repeatability yielded 13.469 hits (conducted on 21.08.2026). The number of publications in this field has grown rapidly since the early 2000s, with over 900 papers from 2025 alone (see supplementary Figure S1).

Despite this enormous interest, approaches for quantifying repeatability vary substantially across studies. A meta-analysis of 759 repeatability estimates from early individuality studies found that the majority of studies measured individuals only twice (i.e. the mode of the distribution), with a mean of 4.41 measures and a range of 2–60 (Bell et al., 2009). Guidelines for repeatability estimation confirm that most studies cluster at small sample sizes and only two repeated measures; a design sufficient to yield precise confidence intervals only when repeatability is very high (Wolak et al., 2012). Methodological choices regarding the statistical framework also vary: correlation-based methods and interclass correlation coefficients derived from mixed-effects models are the most common approaches with Bayesian approaches being less commonly used (MacKinlay & Shaw, 2023). Despite widespread acknowledgment that repeated measures are necessary to estimate among-individual variance, a recent survey of researchers found that 31% reported having used single measurements per individual (Sánchez-Tójar et al., 2022).

These practices sit in tension with the above outlined growing recognition that individuality can fluctuate across ontogeny. As Stamps & Groothuis (2010a) put it: *“[…] a ‘snapshot’ view of personality, which is based on descriptions of behaviour at a single age or life stage, provides an inadequate foundation for studies of personality across ecological and evolutionary scales of time and space“*. Indeed, as the field’s ability to obtain temporally rich, longitudinal behavioral datasets only improves, an added complication to the quantification of repeatability is that repeatability operating at different temporal scales can have very different meanings (Araya-Ajoy et al., 2015; Biro & Stamps, 2015; Mitchell et al., 2020). For example, repeatability of observations collected minutes apart may quantify short-term persistence in behavioral states, whereas repeatability of observations collected across days, seasons, or developmental stages represents longer-term stability of individual differences. Individuals may thus be highly consistent within days but not across development or, conversely, may show high variability within days, but remain somewhat stable in their relative behavioral expression across longer development-length intervals. Thus, repeatability, and its biological interpretation, should be understood as conditional on the temporal scale and developmental window over which it is estimated. In sum, if repeatability estimates are sensitive to when, how often, and over what interval behavior is measured, then comparisons across studies and drawn conclusions may be substantially shaped by methodological choices as much as biology.

### The present study

Here, we leverage high-resolution tracking data to examine how measurement choices affect repeatability estimates across ontogeny. We tracked 45 clonal Amazon mollies (*Poecilia formosa*) at 5 frames per second across the entire first four weeks of their lives—a dataset notable for its temporal resolution and ontogenetic coverage (Ehlman et al., 2023, 2025; Scherer et al., 2023). Because Amazon mollies reproduce clonally, among-individual differences in behavior cannot be attributed to genetic differences, providing a uniquely clean window onto the developmental and stochastic processes that generate behavioral individuality (Laskowski et al., 2022). Using these data, we systematically vary the measurement choices that researchers commonly make, including the level of temporal aggregation (e.g. averaging across minutes vs across hours), the sampling rate (e.g. every minute vs every hour), and resulting number of repeated measures, as well as the interval between measurement end points (e.g. repeatability between the first and seventh day vs the first and twenty-eights day). We then examine their consequences for repeatability estimates and the underlying within- and among-individual variances across developmental stages. Finally, we close with key insights and practical recommendations for researchers studying individuality across development.

## Methods

### Ethics and data availability

All animal care and experimental protocols complied with local and federal laws and guidelines and were approved by the appropriate governing body in Berlin, Germany, the Landesamt fur Gesundheit und Soziales (LaGeSo G-0224/20).

The behavioral data have been made available alongside previous publications (Ehlman et al., 2025; Scherer et al., 2023) and can be found in this Dyrad data repository (https://doi.org/10.5061/dryad.x69p8cztw). Additionally, we have made all analysis code available on the Open Science Framework (https://osf.io/748xz/overview?view_only=14268af6bbd144529961aba4edd90b58).

### Data and experimental design

Full methodological details regarding the study species, housing conditions, feeding regimens, and experimental setup have been reported elsewhere (Ehlman et al., 2025; Scherer et al., 2023). Below, we summarize the key points. Genetically identical gravid Amazon mollies (*Poecilia formosa*) were drawn from a single isogenic stock maintained at Humboldt Universität zu Berlin (Berlin, Germany). The Amazon molly is a gynogenetic freshwater fish species (Lamatsch et al., 2002; Schlupp, 2005) known for a rich behavioral repertoire across development (Bierbach et al., 2017; Laskowski, Bierbach, et al., 2022; Scherer et al., 2023, 2024). Offspring from three of these isogenic females served as the subjects in behavioral experiments. Because all three mothers were genetically identical and sourced from the same stock tank, maternal genetic variation was effectively eliminated; individual maternal identity was nonetheless included as a factor in all statistical models to account for any remaining maternal influences (see below). Together, the three mothers contributed 45 experimental fish, which were transferred to individual observation tanks on the day of birth (supplementary Figure S2). Beginning the following day (i.e. their first complete day of life) fish were recorded from above using a Basler acA5472-5 gm camera equipped with a 16 mm lens (Basler Lens C11-1620-12M-P f16mm) at 5 frames per second for eight continuous hours daily, continuing until day 28. Since Amazon mollies reach sexual maturity at around three months of age (Scherer et al., 2023), this observation window encompassed roughly one-third of their developmental period. Although overhead recording precluded capture of vertical movement, tank water was kept shallow (approximately 7 cm), such that movement was effectively constrained to two dimensions. Fish were maintained on a 12:12 h light–dark cycle at an air temperature of approximately 24 ± 1°C and fed once daily during a two-hour window immediately after the filming period, using a stationary food patch of Sera vipan baby fish food set in agar. Each observation tank was lit from below by four LEDs (each 100 cm long, 12 V, color temperature 5500 K, light output ∼1570 lm). The tanks were constructed from white polyethylene to promote even, diffuse illumination. Bottom lighting reduced surface glare and facilitated automated video tracking using the software Biotracker. This setup produced time series of fish positions in *x*–*y* coordinate space at a temporal resolution of 0.2 seconds.

### Variables

Our outcome of interest is activity. To this end, we use the *x*–*y* coordinates to compute the average step size (in cm) moved per 0.2 second. As independent variables we include fish identity, maternal identity, standard length (i.e. fish length from the tip of the snout to the end of the peduncle, thus excluding tail fin) across the experiment, and where applicable, age in experimental days.

### Repeatability across ontogeny

We compute repeatability to quantify the extent to which individual differences in activity are stable across ontogeny (i.e. indicative of individuality). Repeatability is a ratio of variances: it quantifies variation between individuals relative to total variation (i.e. the sum of between- and within-individual variation). Mixed modeling approaches are commonly used to quantify these variance components. Using R version 4.4.3 (R Core Team, 2023), we compute repeatability in two ways. First, we model how daily activity (i.e. average daily step size using all of the available data) changes across the measurement period using a mixed modeling framework (lme4-package, Bates et al., 2015). This allows us to estimate individual behavioral reaction norm components (i.e. the set of behavioral phenotypes that an individual produces across time; Dingemanse et al., 2010) as well as repeatability and its underlying among - and within-individual variance components across different experimental days (Laskowski, Bierbach, et al., 2022). We use bootstrapped confidence intervals for inference. The response variable is log-transformed for normalization, meaning that our model captures proportional rather than absolute differences step size. The fixed effects include centered age in days, standard length (cm), and maternal identity, and we allow individual-specific trajectories over time by including random intercepts for individuals and random slopes for age. From this model, we extract the variance components of the random intercept (*V*_ind_), random slope (*V*_slope_), their covariance (*Cov*_ind,slope_), and the residual variance (*V*_res_ = *σ*^2^). For a given centered age *H* (i.e. day 1, 7, 14, 21, 28), we derive the among-individual population variance at that age according the following formula:

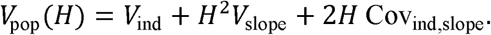

Repeatability of activity on the log scale at age *H* is then

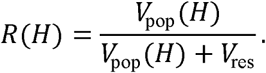

We conduct additional sensitivity analyses to evaluate the robustness of our estimates to assumptions about residual within-individual variation and temporal dependence of measures taken closely in time (i.e. temporal autocorrelation; see supplements Figures S3.1-S3.2). In the analyses described above, residual within-individual variance is assumed to be homogeneous across development, and residual observations are assumed to be temporally independent. However, previous work has shown that these assumptions tend to not hold in densely sampled data and that neglecting these factors can affect repeatability estimates (Cleasby et al., 2015; Mitchell et al., 2020, 2021; O’Dea et al., 2021). We therefore fit additional models that relax these assumptions by allowing within-individual variance to vary across development and by explicitly modeling temporal autocorrelation using autoregressive AR(1) structure (Mitchell et al., 2020). These analyses are implemented in a Bayesian framework using brms (Burkner PC, 2013).

Following the approach outlined above (Laskowski, Bierbach, et al., 2022), we then compare the resulting repeatability estimates across development obtained from a model that does not model changes in within-individual variance or autocorrelation, with estimates from models that explicitly model these quantities. Because these models are more data-intensive, we are unable to fit them to our main analyses which estimate repeatability across different day pairs, aggregation levels, and sampling rates (see below). In such cases, the relevant down-sampled subsets of our data are often insufficient to reliably estimate the additional variance and autocorrelation parameters.

### Emulating different empirical workflows

Second, for our main approach, we use the R-package *rptR* to compute repeatability (Stoffel et al., 2017). We use bootstrapped 95% confidence intervals and permuted *p*-values (*α* = .05) for inference. Here, we compute repeatability at any given day (or between two days) by only including data from these time points (see below). As with the previous model, we log-transformed the outcome for normality. As we typically only include at most two measurement days in our models, we chose a simpler random effects structure and only included a random intercept for individuals, avoiding convergence issues. Additionally, this enabled us to keep a consistent random effects structure across all of our models. The fixed effects included age in days (when there was more than one day) and maternal identity. We removed standard length based on the results of the initial model described above. We inspected model assumptions visually by plotting residuals and q-q plots.

The high-resolution data allow us to emulate several empirical workflows and to explore their impact on repeatability patterns. First, we explored the impact of *aggregation level.* Assuming that researchers had access to all of the available data, we examined how repeatability changes when we aggregate within different intervals, ranging from minutes to days. Second, we explored the impact of *sampling rate*. Assuming that researchers conducted behavioral observations for 10 minutes, we explored different rates at which these 10-minute observation windows were sampled, ranging from continuously (i.e. many repeated measures) to once per day (i.e. few repeated measures). For sampling rates which resulted in only one or two measures per day, we additionally investigated the impact of the *measurement time during the day*. Third, we explored the impact of frame rate. What if instead of recording at 5 frames per second, behavior was recorded, for example, at 1 frame per second or even 1 frame per minute? Fourth, we explored how the inclusion of measures on days occurring between the two measurement days shaped repeatability. To illustrate, suppose that a researcher wants to quantify repeatability between day 1 and 7. How does the repeatability estimate change when we only include measures from those two days vs measures from all the intermediate days as well. These different workflows reflect measurement choices and constraints that researchers may make and face, showcasing their potential influence on repeatability estimates.

## Results

To properly interpret the results, it is important to keep in mind that we log-transformed the outcome, meaning that the estimated variances are also on log scale. Log-scale variances should be interpreted in terms of proportional rather than absolute differences in activity. Specifically, an among-individual variance of *V*_ind_ on the log scale implies that individuals differ by a factor of approximately 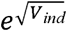 in their average step size, that is, the standard deviation on the log scale corresponds to a multiplicative spread around the mean on the raw scale. For example, an among-individual standard deviation of 0.5 log-units means that individuals one *SD* apart in the population differ by a factor of roughly *e*^0.5^ ≈ 1.65, or about 65%, in their activity level. Similarly, the residual variance captures within-individual moment-to-moment fluctuations expressed as proportional deviations from an individual’s own mean. Resulting repeatability estimates should therefore be interpreted as indicating how consistent individuals are in their relative (proportional) activity level across time: a high repeatability means that an individual that is, say, twice as active as the average on one occasion tends to remain twice as active on a later occasion.

### Variance components across ontogeny

First, we modelled daily average step size across ontogeny, adjusting for maternal identity and standard length, and including random intercepts and random slopes for experimental day per individual fish (Figure 1, top panel). Activity declines significantly over the experimental period (*β_day_* = −0.049, 95% CI [−0.061, −0.036]), indicating that on average, fish become proportionally less active as they age. Specifically, the model shows that, on average, fish are moving approximately 1 − *e*^−0.049^ = 4.8% less per additional day. Maternal identity shows some evidence of an effect, with offspring of mother ‘m4’ being more active than the reference mother (*β* = 0.318, 95% CI [0.056, 0.573]), while the confidence interval for mother ‘m2’ just included zero (*β* = 0.450, 95% CI [−0.005, 0.887]). Standard length shows a positive but non-significant association with activity, indicated by the confidence interval including zero (*β* = 0.178, 95% CI [−0.051, 0.399]). Because including standard length does not alter the substantive conclusions, we omit it from subsequent analyses for parsimony. In the supplements we show that our main results are qualitatively identical when we include standard length (supplementary Figures S21-S24).

**Figure 1:**
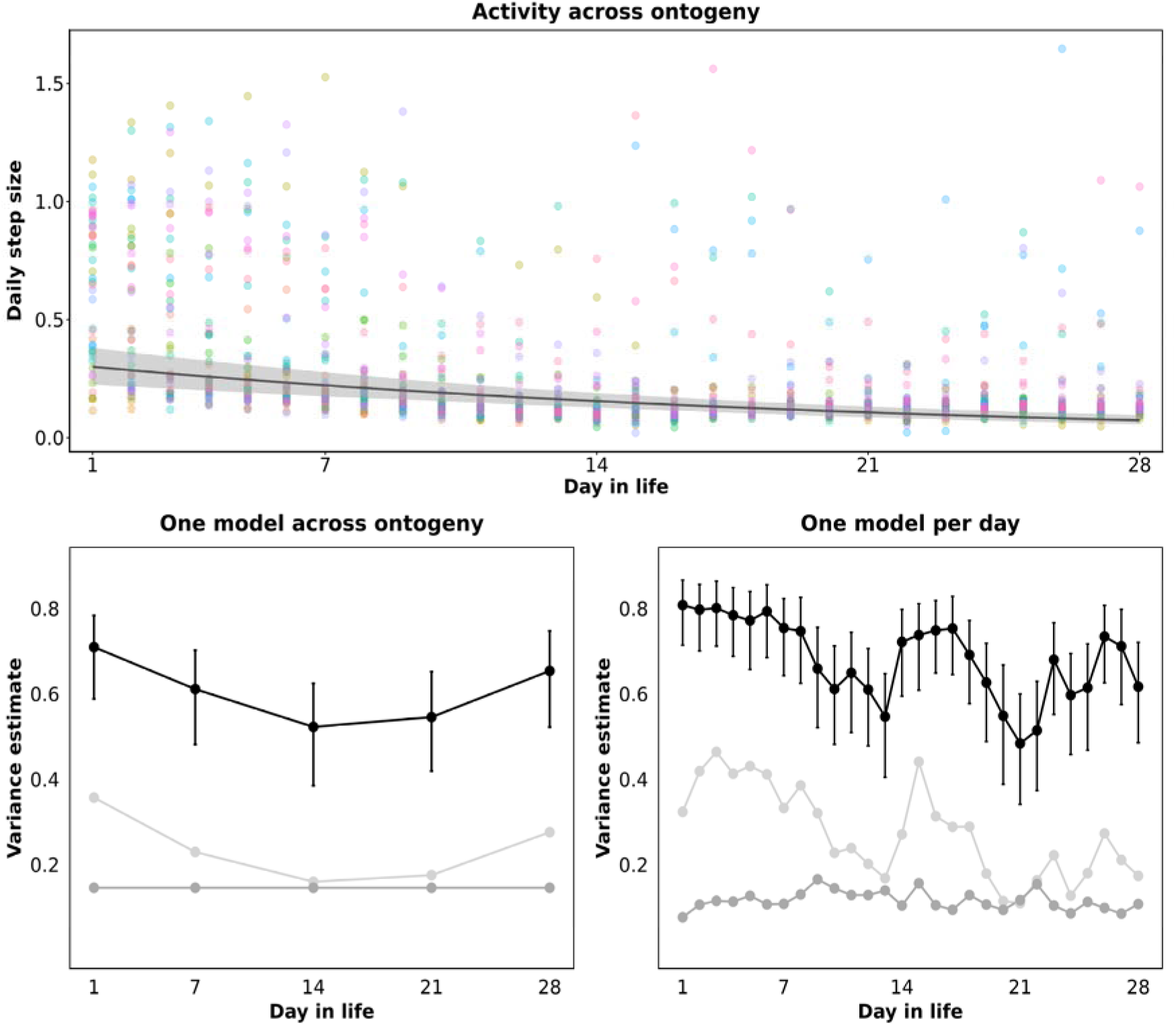
Activity and variances across ontogeny. Across all panels, the horizontal axis indicates the experimental day, corresponding fish age. The top panel shows the model-estimated changes in activity (vertical axis) across age, with 95% confidence intervals. To facilitate interpretation, the estimated age effect is shown on the raw activity scale. The scatterplot shows raw daily activity data for all fish. The bottom left panel shows variance components (on log-activity scale) and repeatability across ontogeny (vertical axis), estimated from the model shown in the top panel. The bottom right panel shows variance components (on log-activity scale) and repeatability across ontogeny (vertical axis), estimated by fitting separate models to hourly activity data for each day.

The random effects structure reveals both individual differences in activity and individual differences in plasticity. The substantial among-individual variance in intercepts (*SD_ind_* = 0.398, 95% CI [0.309, 0.489]) indicates that individuals consistently differ in their average activity level, that is, fish differ in the elevation of their behavioral reaction norm at mean age (Dingemanse et al., 2010). The among-individual variance in slopes (*SD_slope_* = 0.029, 95% CI [0.023, 0.036]), though smaller in magnitude, indicates that individuals also differ in how their activity changes across ontogeny, that is, fish differ in the slope of their behavioral reaction norm, reflecting individual differences in developmental plasticity. The intercept– slope correlation was weakly negative (*r* = −0.14, 95% CI [−0.445, 0.206]) but nonsignificant, suggesting no association between mean activity levels and changes in activity across ontogeny. Together, these results justify the age-specific repeatability approach: because individuals differ in both elevation and slope, the degree to which individual differences are consistent depends on the developmental stage at which behavior is measured.

Following Laskowski et al. (2022), we use this model to estimate among- and within-individual variances, as well as repeatability at different ontogenetic time points (Figure 1, bottom left panel). We find that repeatability changes as a function of age, largely tracking changes in among-individual variance. Estimated repeatability is relatively high early in ontogeny, decreases across the first two weeks of life, and increases again during the latter half of the experiment. However, we also note that the confidence intervals of estimates across time are overlapping. We contrast these estimates with repeatability estimates computed from separate models fitted to each day (Figure 1, bottom right panel). Each model uses all available data from a single day. Activity was aggregated within each hour of the day, yielding one mean activity value per individual per hour. These day-specific models broadly reproduce the pattern observed in the model using data spanning the entire measurement period: repeatability first decreases before increasing again, and estimates track among-individual variances. However, modelling each day separately also reveals a temporary increase in repeatability during the third week of life. Thus, repeatability does not appear to change continuously or smoothly across ontogeny. Consequently, conclusions about developmental changes in repeatability may depend on the particular developmental window sampled.

Sensitivity analyses incorporating heterogeneous within-individual variance or autocorrelation produced qualitatively different results depending on aggregation level. With daily aggregation, within-individual variance declined across development (*β_sigma_* = −0.048, 95% credible interval [−0.056, −0.041]), driving a corresponding increase in repeatability (supplementary Figure S3.1). Modeling autocorrelation at this level reproduced the u-shaped repeatability pattern seen in the main analyses but at lower overall magnitude. Additionally, the relatively wide confidence intervals suggest that the daily activity data provide insufficient repeated measures to precisely estimate variance components alongside autocorrelation. And, indeed, fitting both quantities simultaneously was not feasible at this aggregation level.

With hourly aggregation (supplementary Figure S3.2), the same sensitivity analyses produced less dramatic departures from the main results. Within-individual variance still declined with age, but the slope was shallower (*β_sigma_* = −0.022, 95% credible interval [−0.025, −0.020]) and the early-ontogeny elevation in within-individual variance relative to among-individual variance was less pronounced. Accordingly, repeatability estimates were only modestly reduced relative to the homogeneous-variance model. Modeling autocorrelation across hours within the same day preserved the u-shaped pattern and, unlike at the daily level, actually increased repeatability estimates relative to the model without autocorrelation. Here, the autocorrelation absorbed within-individual variance while leaving among-individual variance intact. Resulting repeatability patterns are similar to those obtained from daily data (Figure 1, bottom left panel), implying that modeling autocorrelation in hourly data has a similar effect as averaging data across hours (supplements Figures S3.1-S3.2). When both heterogeneous variance and autocorrelation were modeled jointly, within-individual variance showed only a shallow decline (*β_sigma_* = −0.008, 95% credible interval [−0.010, −0.006]), and the u-shaped repeatability pattern was retained.

### Emulating different empirical workflows

#### 1. Repeatability across ontogeny at different aggregation levels

We find that repeatability estimates increase at higher aggregation levels, where each data point is the result of averaging across more observations and longer measurement windows (Figure 2). The largest increase occurs between the minute-level and hourly aggregation levels. These patterns are consistent with mathematical theory and simulations demonstrating that repeatability of means is generally higher than repeatability of unaggregated data because averaging reduces measurement error or true within-individual variation (Nakagawa & Schielzeth, 2010). However, when repeatability is computed from only two observations per individual at the daily aggregation level, estimates sometimes decrease. These estimates are based on comparatively little information and are therefore likely more sensitive to measurement uncertainty. Comparing Figure 2 with the equivalent figure including all intermediate days, we see the same pattern of increasing repeatability with aggregation level (supplementary Figure S5). The results from models including intermediate days are smoother and do not show the late decline apparent in Figure 2, likely because more data are available to estimate repeatability more precisely. Additionally, repeatability estimates tend to be higher when measurement days are closer in time, that is, at lower day differences, and earlier in ontogeny. As in Figure 1, repeatability does not decline gradually across ontogeny but instead increases again towards the middle and end of the experiment.

**Figure 2:**
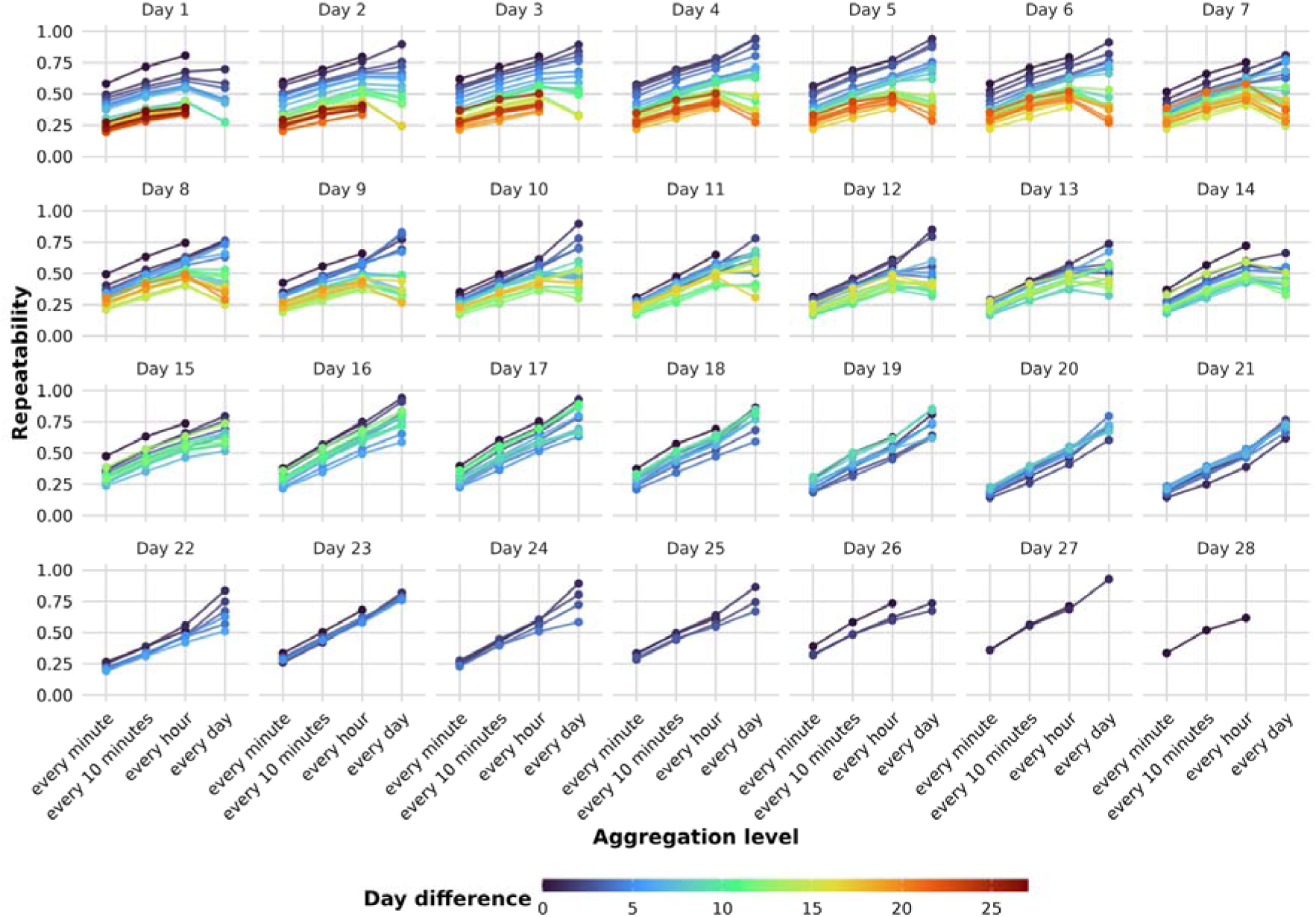
Repeatability across aggregation levels. Each panel represents a focal day in the fish’s life. The vertical axis shows statistically significant (*α* < .05) repeatability estimates. Each filled circle corresponds to one model fitted to data from that focal day and, except where the day difference was zero, some subsequent day (separated from a focal day by the indicated day difference). Colors of circles and connecting lines indicate the number of days separating the two measurement days (day difference; 0–27 days). The horizontal axis indicates the temporal aggregation level applied to the activity data: every minute, every 10 minutes, every hour, or every day. For example, at the *every minute* level, raw activity observations (recorded at 5 frames per second) were averaged within each minute, yielding one observation per individual per minute. A day difference of zero denotes estimates based on data from the focal day alone. Repeatability could not be estimated at the daily aggregation level for these models because this aggregation yields only one observation per individual per day. Other missing estimates indicate that the repeatability estimate did not reach the statistical significance threshold.

For the same models, Figure 3 and supplementary Figure S4 (including intermediate days) show heatmaps of repeatability estimates and the underlying among- and within-individual variances. Among-individual variances are qualitatively similar across aggregation levels. However, we observe a relatively large reduction in within-individual variance between the minute-level and hourly aggregation levels, explaining the relatively steep initial increase in repeatability estimates. At the daily aggregation level, estimates are less often statistically significant when the interval between measurement days is large. This should not be interpreted as evidence that repeatability is absent; rather, it indicates that the available data do not provide sufficient evidence to reject the null hypothesis of zero repeatability at the chosen significance threshold.

**Figure 3:**
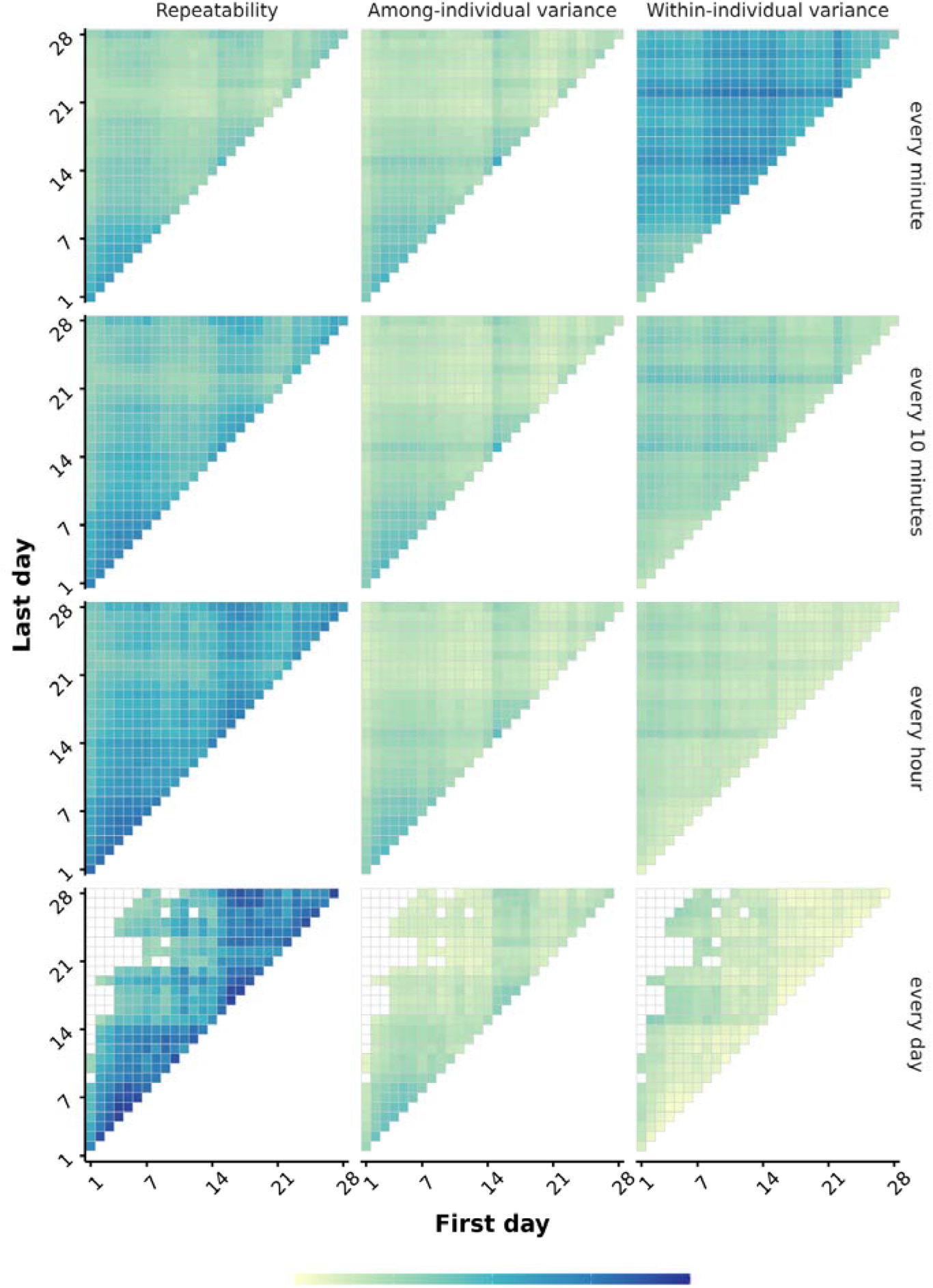
Repeatability and variance components across aggregation levels. Rows represent the temporal aggregation level applied to the activity data: every minute, every 10 minutes, every hour, and every day. For example, at the every minute level, raw activity observations (recorded at 5 frames per second) were averaged within each minute, yielding one observation per individual per minute. Columns show, from left to right, heatmaps illustrating repeatability (*R*), among-individual variance, and within-individual variance estimates; variance components are reported on the log-activity scale. Each cell within a heatmap represents a significant estimate (*α* < .05) from a separate model fitted to data from the pair of days indicated by the axes. The horizontal axis gives the first day and the vertical axis gives the second day; only unique day pairs are shown. Cells on the diagonal therefore represent models fitted to observations from a single day, whereas off-diagonal cells represent models fitted to observations from two different days. Cell color denotes the estimated value. At the daily aggregation level, estimates from same-day models cannot be obtained because each individual contributes only one observation per day at this aggregation level. Empty cells elsewhere indicate estimates that did not reach the statistical significance threshold. Repeatability is bounded between 0 and 1. Variance components are not inherently bounded, but all estimated values in this dataset were below 1; therefore, a common color scale is used across panels.

#### 2. Repeatability across ontogeny at different sampling rates

We find that repeatability estimates decrease with lower sampling rates of 10-minute observation windows (Figure 4). As the number of available observations drops to two or one per day, repeatability estimates sometimes increase. However, estimates at these lowest sampling rates are also highly variable and do not appear to follow a consistent pattern.

**Figure 4:**
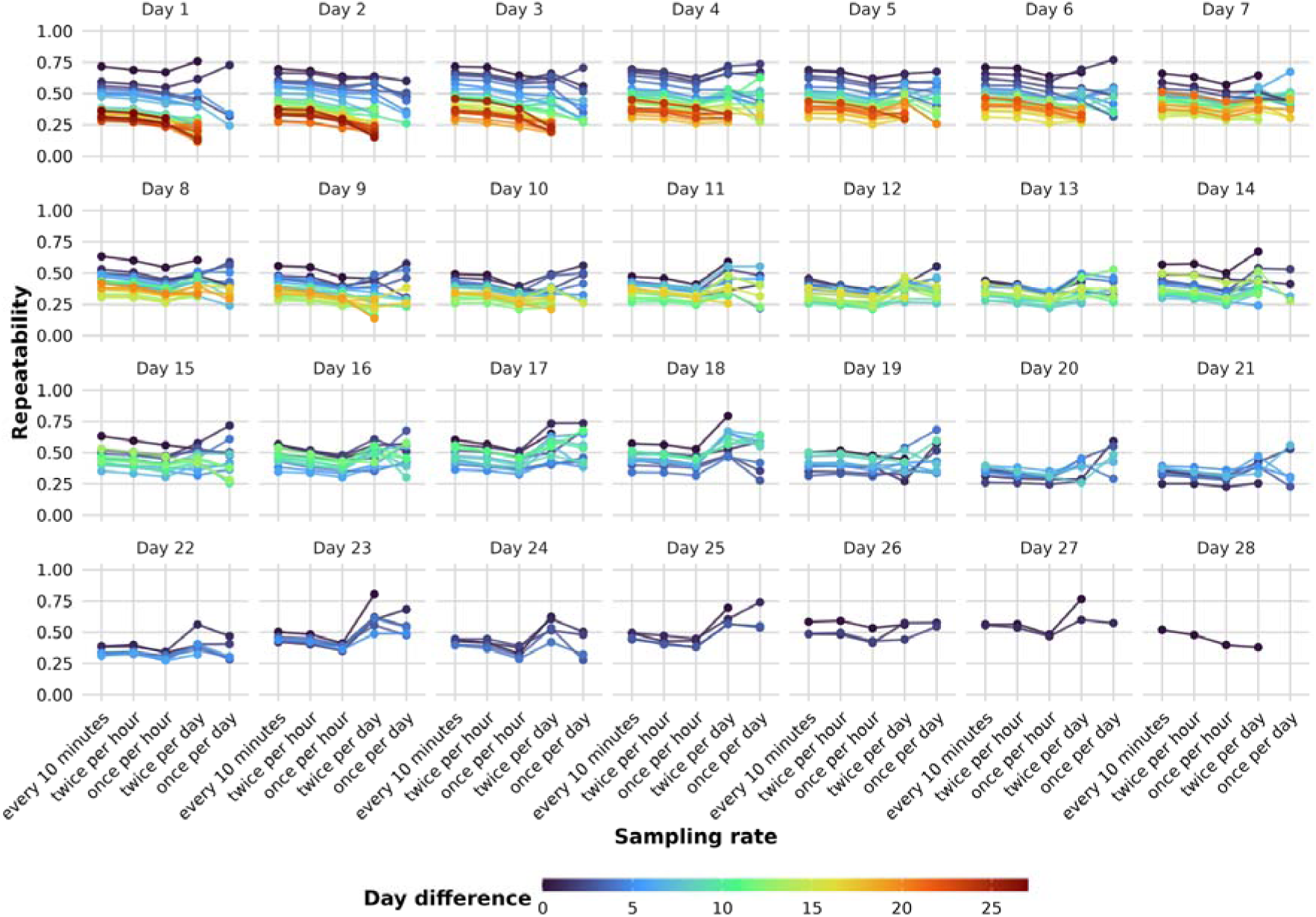
Repeatability across sampling rates. Each panel represents a focal day in the fish’s life. The vertical axis shows statistically significant (*α* < .05) repeatability estimates. Each filled circle corresponds to one model fitted to data from that focal day and, except where the day difference was zero, some subsequent day (separated from a focal day by the indicated day difference). Colors of circles and connecting lines indicate the number of days separating the two measurement days (day difference; 0–27 days). The raw activity observations (recorded at 5 frames per second) were first aggregated across 10-minute observation intervals, resulting in one activity data point per every 10 minutes, per fish. The horizontal axis indicates the sampling rate of these 10-minute observation intervals: continuously every 10 minutes, twice per hour, once per hour, twice per day, and once per day. For example, the *twice per hour* sampling rate means that the model included aggregated activity data from two 10-minute observation intervals per hour, per individual. A day difference of zero denotes estimates based on data from the focal day alone. Repeatability could not be estimated at the once per day sampling rate for these models because this rate yields only one observation per individual, per day. Other missing estimates indicate that the repeatability estimate did not reach the statistical significance threshold.

This greater variability is consistent with increased measurement uncertainty at lower sampling rates, as indicated by the wider confidence intervals at these sampling rates (Figure 5). Results including intermediate days reveal the same overall decreasing trend in repeatability estimates (supplementary Figure S7). As in the previous comparison between analyses with and without intermediate days, including more days produces smoother patterns across day differences and fewer abrupt changes when few observations were available.

**Figure 5:**
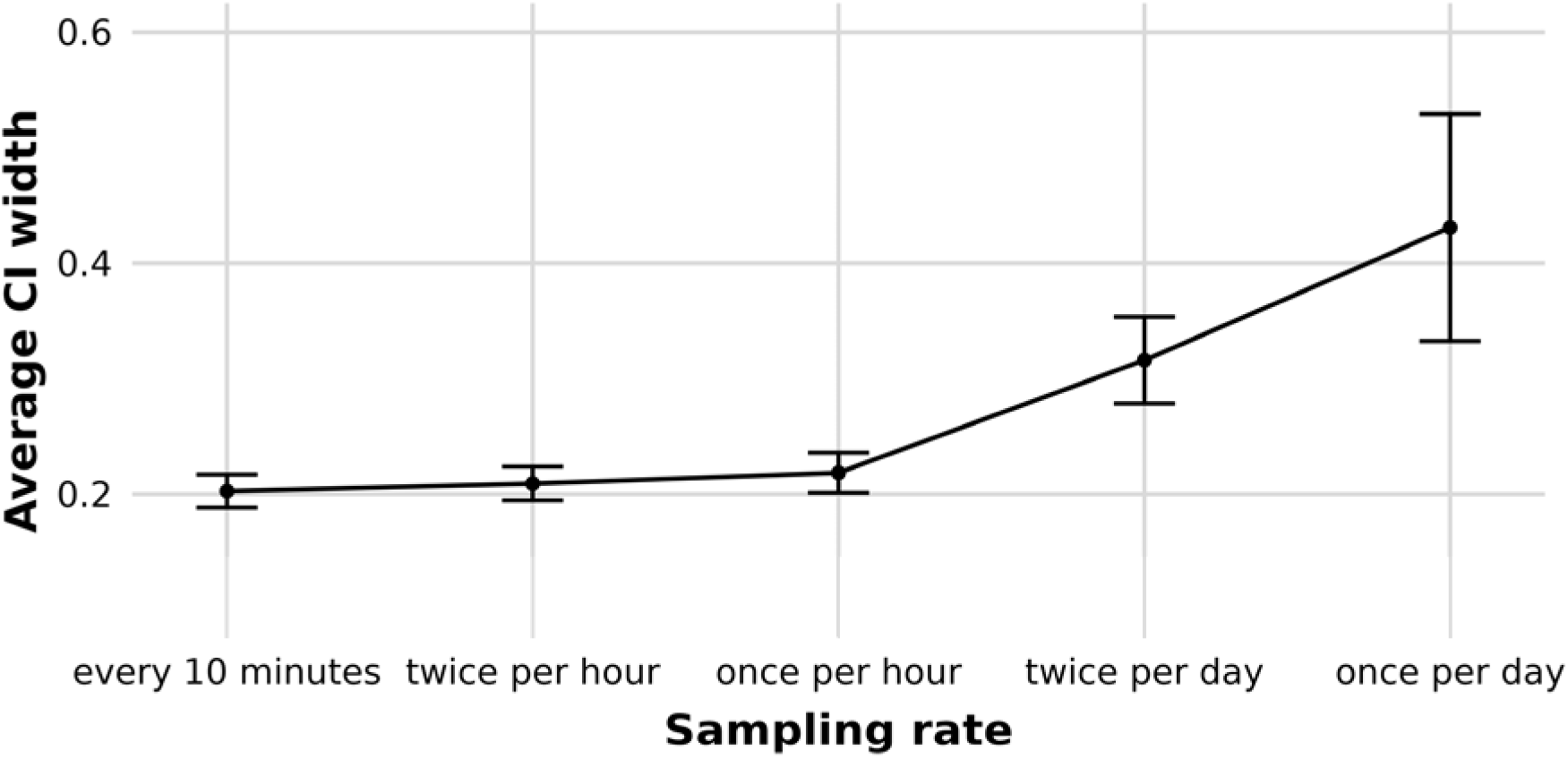
Confidence intervals around repeatability estimates across sampling rates. 95% confidence intervals are based on models fitted to activity data resulting from different sampling rates of 10-minute observation intervals (see Figure 4). The horizontal axis shows these different sampling rates. The vertical axis shows the average width of the confidence intervals across all models computed from data at a specific sampling rate. The error bar indicates the standard deviation around the average width.

For low sampling rates, we additionally explored the impact of measurement time within the day (Figure 6). When only one or two 10-minute observation windows per fish per day are used, repeatability estimates vary depending on when measurements are taken. When two observations per day are obtained from the beginning and end of the day, repeatability estimates are consistently low. In contrast, when observations are obtained from the second or third hour and the corresponding penultimate hours of the day, estimates are higher and more similar to those obtained at denser sampling rates.

**Figure 6:**
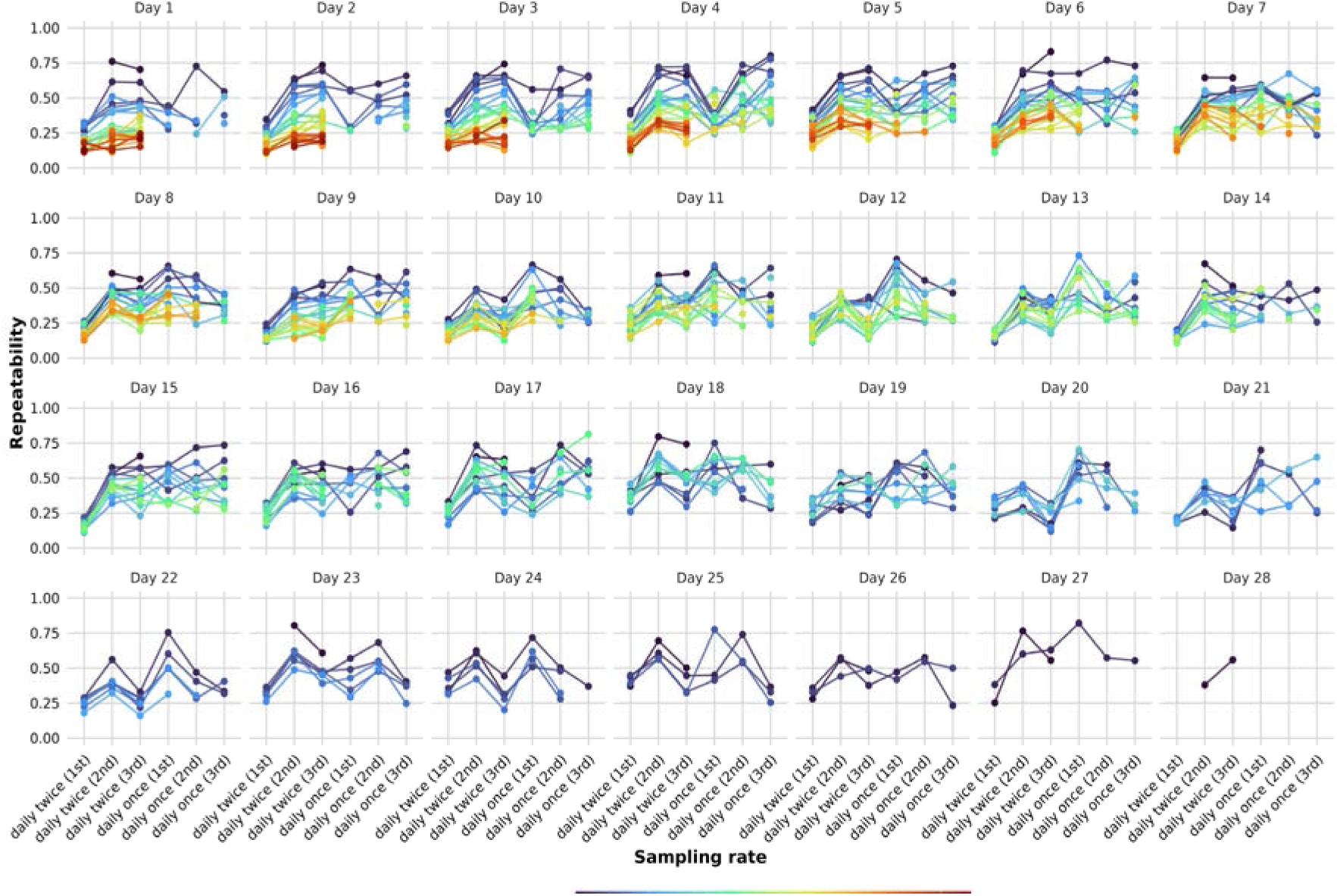
Repeatability across sampling rates and measurement times. Each panel represents a focal day in the fish’s life. The vertical axis shows statistically significant (*α* < .05) repeatability estimates. Each filled circle corresponds to one model fitted to data from that focal day and, except where the day difference was zero, some subsequent day (separated from a focal day by the indicated day difference). Colors of circles and connecting lines indicate the number of days separating the two measurement days (day difference; 0–27 days). The raw activity observations (recorded at 5 frames per second) were first aggregated across 10-minute observation intervals, resulting in one activity data point per every 10 minutes, per fish. The horizontal axis indicates the sampling rate of these 10-minute observation intervals: twice per day using the first and last 10 minutes of each day, twice per day using the second and second-to-last 10 minutes of each day, twice per day using the third and third-to-last 10 minutes of each day, once per day using the first 10 minutes of each day, once per day using the second 10 minutes of each day, and once per day using the third 10 minutes of each day. A day difference of zero denotes estimates based on data from the focal day alone. Repeatability could not be estimated at the once per day sampling rate for these models because this rate yields only one observation per individual, per day. Other missing estimates indicate that the repeatability estimate did not reach the statistical significance threshold.

The variance decomposition (supplementary Figure S10) indicates lower among-individual variance and higher within-individual variance when observations are obtained from the beginning and end of the day. This pattern may reflect systematic differences in fish behavior at these times of day, ‘artefacts’ introduced by the tracking software or hardware at the boundaries of the recording period (e.g. fish anticipating the red light turning on at the start and off at the end of the recording period), or both. When only a single observation per fish per day is used, similar patterns are sometimes observed, but less consistently. For observations collected during the first 10 minutes of each day, both variance components were small in magnitude, suggesting limited absolute variation among and within individuals during this period. When both components are small, modest absolute differences in either variance component can produce large differences in repeatability because repeatability is calculated as a ratio of variance components. This explains the large variability in repeatability estimates across different measurement times of the same day.

Overall, repeatability estimates based on less extreme measurement times within the day are more similar to one another. However, estimates can still fluctuate substantially, highlighting that repeatability cannot be estimated reliably from only one or two measurements per fish per day in this dataset.

#### 3. Impact of frame rate

We find that a frame rate of 1 frame per second yields results that are indistinguishable from those computed using observations tracked at 5 frames per second (supplementary Figures S11-S15). However, when the frame rate is further reduced to 1 frame per minute, repeatability estimates are substantially lower, in some cases decreasing by approximately 0.5 repeatability units (supplementary Figures S16-S20).

Although qualitative patterns are largely preserved at 1 frame per minute, it becomes more difficult to distinguish differences in repeatability between developmental stages. The variance plots identify low among-individual variance and high within-individual variance as the primary contributors to these patterns. However, as observations are aggregated across more data points (supplementary Figure S16), repeatability estimates approach those observed at higher frame rates.

#### 4. Impact of coverage and intermediate days

Lastly, Figure 7 shows that the temporal placement of intermediate measurement days can matter more than whether intermediate days are included at all. Suppose that a researcher wishes to estimate repeatability across the first four weeks of life. Rather than including only days 1 and 28, the researcher decides to include intermediate days but is not be able to record fish every day. Which days should they include?

**Figure 7:**
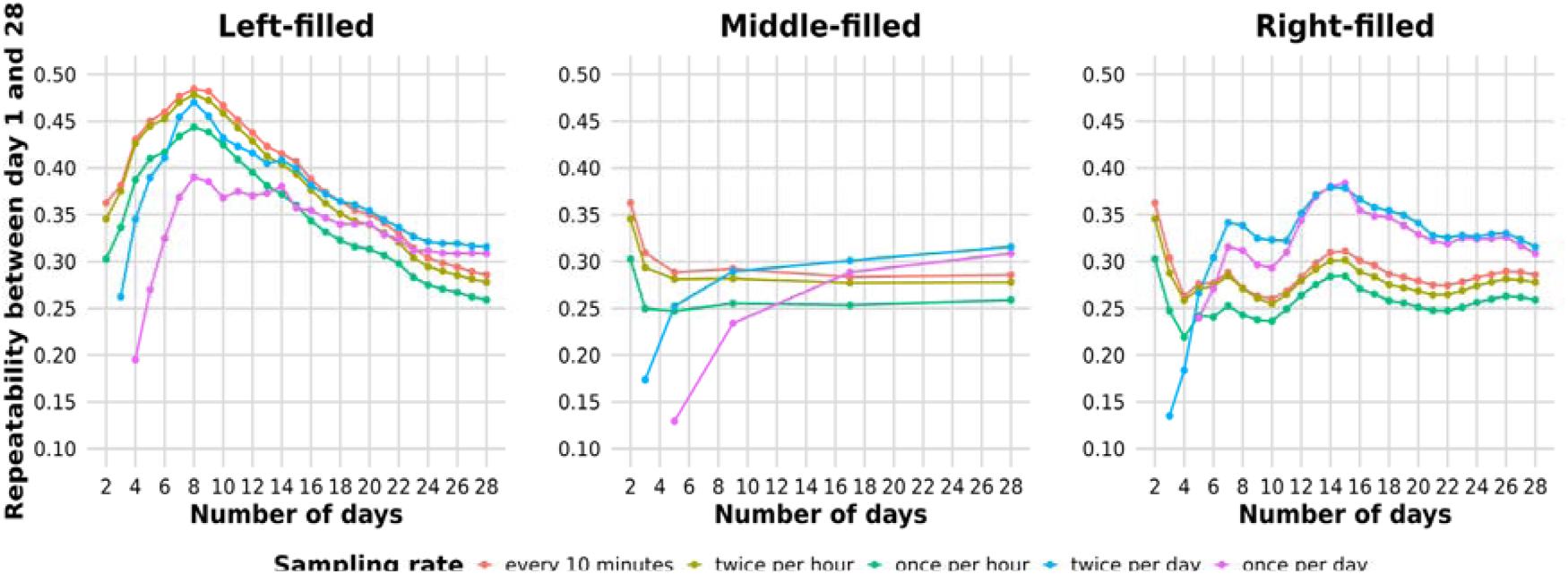
Repeatability between day 1 and 28 across sampling rates and number of included days. Within a panel the vertical axis shows repeatability estimates (*R*) between activity measures on day 1 and 28. The horizontal axis indicates the total number of measurement days included in the model, ranging from 2 (days 1 and 28 only) to 28 (all measurement days). The vertical axis shows statistically significant (*α* < .05) repeatability estimates. Panels represent alternative strategies for adding the intermediate measurement days. In the *left-filled* approach, days were added sequentially from the beginning of the study period; in the *right-filled* approach, they were added sequentially from the end. In the *middle-filled* approach, additional days were distributed as evenly as possible across the interval: starting with days 1 and 28, day 14 was added first, followed by the midpoints of the resulting intervals (e.g. days between 1–14 and 14–28), and so forth. Missing estimates indicate that the repeatability estimate did not reach the statistical significance threshold.

When comparing different strategies for adding intermediate days, repeatability estimates fluctuate depending on whether data are added from earlier or later developmental stages. This pattern is consistent with the preceding analyses, which showed that repeatability tended to be higher earlier and lower later in development. These age-specific differences are reflected in repeatability estimates that include days from different developmental stages.

In contrast, when intermediate days are added from the middle of the study period, estimates are more stable. Figure 7 also shows that repeatability estimates based only on the first and last day tend to be higher than estimates obtained when intermediate days are included. Thus, in sparse longitudinal designs, repeatability estimates depend not only on the number of measurement days but also on their placement across ontogeny.

## Discussion

Our results show that repeatability estimates of locomotor activity in clonal Amazon mollies are sensitive to a range of measurement choices that researchers routinely make. These findings suggest that a portion of the variation in repeatability estimates reported across the individuality literature may reflect methodological heterogeneity as much as genuine biological differences between species, populations, or traits. However, we also need to acknowledge there is no single ground truth for repeatability. Even with behavioral data recorded at 5 frames per second throughout the first 28 days of life, what constitutes a biologically meaningful estimate of repeatability ultimately depends on the question being asked. Thus, some of the variation in repeatability estimates highlighted here may reflect differences in the temporal scales at which underlying biological processes operate.

### Repeatability across developmental stages

Across all measurement conditions, locomotor activity was repeatable, adding to a growing literature documenting individual differences in the absence of genetic or environmental variation (Bierbach et al., 2017; de Bivort et al., 2022; Freund et al., 2013, 2015; Laskowski, Bierbach, et al., 2022; Scherer et al., 2023). However, repeatability was not uniform across ontogeny: it declined over the first two weeks of life before recovering in the latter half of the experiment. This pattern was driven primarily by changes in among-individual variance rather than within-individual variance: individuals differed in both the elevation and slope of their activity trajectories (Dingemanse et al., 2010). As a result, among-individual differences, and hence repeatability, depended on the developmental stage at which behavior was assessed.

The early-life decline in repeatability echoes the weak evidence for higher repeatability in juvenile compared to adult ectotherms reported in the to-date largest meta-analysis of repeatability estimates (Bell et al., 2009). This pattern may reflect an initial consolidation of individual behavioral trajectories before within-individual plasticity begins to diversify them (Ehlman et al., 2025; Walasek, Panchanathan, et al., 2025). The subsequent recovery later in the experiment may indicate renewed canalization as developmental trajectories stabilize, potentially analogous to increasing behavioral consistency observed across human adulthood (Roberts & DelVecchio, 2000). In a clonal system, where genetic variation is absent, individual differences in behavioral reaction norm slopes represent a direct expression of developmental plasticity (Stamps & Groothuis, 2010a, 2010b). The observed negative intercept–slope correlation, though uncertain, further hints that more active individuals may show less developmental change, consistent with a pace-of-life perspective in which highly active individuals canalize their behavior earlier (Réale et al., 2007).

### Aggregation effect

We observe pronounced effects of temporal aggregation: as observations were averaged across longer windows, from minute-level to hourly to daily aggregation, repeatability increased. This pattern is consistent with the mathematical relationship between aggregation and repeatability: because within-individual variance is reduced by averaging while among-individual variance is preserved, repeatability of means is always higher than repeatability of raw observations (Nakagawa & Schielzeth, 2010). Researchers should therefore be explicit about aggregation level when reporting repeatability estimates, and comparisons across studies, including systematic meta-analyses, should account for this source of variation.

### Number of measures and measurement uncertainty

Consistent with theoretical predictions (Wolak et al., 2012), we found that lower sampling rates, corresponding to fewer observations per individual per day, produced wider and more variable confidence intervals around repeatability estimates. To-date, a large proportion of published repeatability estimates are based on only two repeated measures per individual (Bell et al., 2009), a design that is sufficient to yield precise confidence intervals only when repeatability is very high (ICC ≥ 0.9) (Wolak et al., 2012). For behavioral traits, which typically show repeatabilities well below this threshold (mean ICC ≈ 0.37; (Bell et al., 2009), two measures per individual are likely to produce estimates with confidence intervals so wide as to render cross-study comparisons unreliable.

A central issue surrounding the number of repeated measures (*k*) is their trade-off with the number of individuals (*n*). Wolak et al. (2012) show that for a given total sampling effort (*n*k*), there is an optimal *k* that minimizes confidence interval width, and that this optimum declines as repeatability increases. For low repeatabilities, as are common in behavioral studies, more measures per individual are needed to precisely estimate within-individual variance. Bell et al. (2009) recommended prioritizing *n* over *k* on the grounds that repeatability estimates did not depend on *k* in their meta-analysis. But Wolak et al. (2012) clarify that while the *point estimate* of repeatability may be unaffected by *k*, the *precision* of that estimate is not. Our results reinforce this distinction: at the lowest sampling rates, repeatability estimates were not only more variable but also more susceptible to artifacts introduced by the time of day at which the sparse observations were collected.

The effective number of repeated observations may also be smaller than the nominal number of observations when measurements are highly temporally autocorrelated—often a property of densely sampled longitudinal datasets. Mitchell et al. (2020) therefore caution that dense sampling alone does not guarantee an unbiased estimate of repeatability and thus recommend either explicitly modeling temporal autocorrelation or separating intensely sampled ‘bursts’ with longer intervals of no measurements.

### Time of day and within-day variation

Our results indicate sensitivity of repeatability estimates to the time of day at which observations are obtained when only one or two measurement windows per individual per day were used. Observations collected at the very beginning or end of the recording period yielded systematically lower repeatability estimates, driven by lower among-individual variance and higher within-individual variance during these periods. This likely reflects a combination of behavioral patterns tied to the light–dark cycle and animals potentially anticipating the tracking start and end when the light turns on and off at the boundaries of the recording session. This implies that when behavioral observations are sparse, the choice of *when* to observe can matter as much as *how often*. Researchers using few observation windows should consider piloting their protocol across different times of day before committing to a fixed schedule, and should report the timing of observations alongside their repeatability estimates.

### The placement of intermediate measurement days

Our analysis of intermediate measurement days revealed that the placement of additional observations across ontogeny can matter more than their number. Including days from earlier developmental stages, when repeatability was higher, inflated estimates relative to including days from later stages. Including days distributed across the middle of the study period produced the most stable estimates. This effect is particularly relevant for studies that aim to characterize repeatability across a broad developmental span but can only afford sparse longitudinal sampling.

### Practical recommendations

Drawing on our analyses, we offer the following recommendations for researchers studying individuality across development:

1. **Report aggregation level explicitly.** Repeatability of aggregated means is systematically higher than repeatability of raw observations. Comparisons across studies are only meaningful when aggregation levels are matched or explicitly accounted for.
2. **Increase the number of measures per individual, particularly when repeatability is expected to be low.** Two measures per individual is insufficient for precise estimation of behavioral repeatability in most systems. Under fixed total effort, the optimal balance between the number of individuals and the number of repeated measures depends on the expected repeatability; Wolak et al.’s (2012) guidelines and the accompanying ICC R package provide a practical tool for a priori power calculations.
3. **Measurement time can affect outcomes.** When only one or two observations per individual per day are feasible, the time of day at which they are collected can distort estimates. Thus, it is advisable to explicitly control for this in the analyses and to conduct robustness checks with other measurement times.
4. **Account for developmental stage.** Repeatability estimates are not interchangeable across ontogeny. Studies that measure behavior at a single life stage should be cautious about generalizing to other stages, and longitudinal designs should explicitly model age-dependent changes in variance components rather than treating repeatability as a constant quantity.
5. **Consider the placement of intermediate measurement days carefully.** In sparse longitudinal designs, adding intermediate observations shifts estimates toward the repeatability of the sampled developmental window. Evenly distributed intermediate days produce more stable estimates of repeatability across the full interval of interest.
6. **Frame rate may be reduced without loss.** In the dataset described here, recording at 1 frame per second produced results indistinguishable from 5 frames per second. Reducing frame rate substantially (i.e. to 1 frame per minute) did degrade estimates, but aggregating across more observations at low frame rates largely recovered the original patterns. This suggests that, in many cases, researchers facing hardware or storage constraints can safely reduce frame rate without major consequences for repeatability estimation, provided sufficient within-day observations are retained.

### Limitations and future directions

We discuss four limitations. First, our system is unusual in its genetic uniformity: because Amazon mollies reproduce clonally, we cannot assess how genetic variation interacts with the measurement factors we examined. However, as our results generally appear to be in line with earlier meta-analyses (Bell et al., 2009) and theoretical work (Nakagawa & Schielzeth, 2010; Wolak et al., 2012), we believe that the measurement choices we investigated here likely have similar effects in genetically diverse populations. Second, our analyses are restricted to a single behavioral trait, i.e. locomotor activity, measured under highly standardized conditions. Repeatability of other traits, particularly those assessed through discrete behavioral assays (e.g. boldness in a novel object test), may respond differently to the measurement factors we examined, and whether these assays are conducted in the laboratory or field, as well as other contextual factors, such as nutrition, may further complicate comparisons (Biro, 2013; Niemelä & Dingemanse, 2017; Walasek, Jovicic, et al., 2025).

Third, we did not examine the consistency of the same behavior across different environmental contexts (e.g. predators present or absent); nor did we examine the consistency of correlations of different behavioral traits (e.g. activity and mating behavior). Extending our work in this way would allow us to gain insight into how measurement choices shape behavioral syndromes, i.e. correlated suites of behaviors that are consistent across situations and/or over time (Sih et al., 2004).

Fourth, for most analyses described above, residual within-individual variance in behavior is modeled as homogeneous across development. This effectively means that changes in repeatability through development are represented in these models as changes in the degree of among-individual variance, with within-individual variance being assumed stable over development (e.g. Figure 1, bottom left panel). In line with previous work, our sensitivity analyses show that allowing for residual variance to vary among individuals as a function of age can impact repeatability estimates (Cleasby et al., 2015; O’Dea et al., 2021). This is consistent with the view that intraindividual variation in behavior—often a measure of individual unpredictability in behavior—can be a biologically important dimension of among-individual differences (Mitchell et al., 2021; Stamps et al., 2012; Westneat et al., 2015). Likewise, explicitly modeling residual autocorrelation reduced the magnitude of repeatability estimates while preserving their developmental pattern (Mitchell et al., 2020). However, these sensitivity analyses had a stronger qualitative impact on daily than on hourly data. Results from hourly data were broadly consistent across models regardless of whether heterogeneous within-individual variance, autocorrelation, both, or neither were included.

Additionally, results were also consistent with the one-model-per-day analyses, which implicitly allow within-individual variance to differ across days (Figure 1, bottom right panel). Across all these approaches, residual variance showed little systematic change through development. This convergence may reflect a limitation of the daily data: 28 repeated measures per individual may be insufficient to precisely partition varying between- and within-individual variances from autocorrelation simultaneously.

Finally, our results speak to the broader challenge of integrating individuality research into conservation science. MacKinlay & Shaw (2023) found that although personality research is increasingly applied in conservation contexts, repeatability was quantified in only half of the reviewed reports. The measurement practices we have explored, including sparse sampling, single life stages, varying aggregation levels, are likely to be even more prevalent in applied settings where logistical constraints are greater. The recommendations we offer are intended to be practically implementable and to help bridge the gap between methodological best practice and field reality.

## References

Adaniya, C. H., Wellman, C. L., Demas, G. E., & Cusick, J. A. (2021). The ontogeny of personality: repeatability of social and escape behaviors across developmental stages in Siberian hamsters (Phodopus sungorus). Journal of Experimental Zoology Part A: Ecological and Integrative Physiology, 335(6). 10.1002/jez.2499

Araya-Ajoy, Y. G., Mathot, K. J., & Dingemanse, N. J. (2015). An approach to estimate short-term, long-term and reaction norm repeatability. Methods in Ecology and Evolution, 6(12), 1462–1473. 10.1111/2041-210X.12430

Bates, D., Mächler, M., Bolker, B., & Walker, S. (2015). Fitting linear mixed-effects models using lme4. Journal of Statistical Software, 67(1), 1–48. 10.18637/jss.v067.i01

Bell, A. M., Hankison, S. J., & Laskowski, K. L. (2009). The repeatability of behaviour: a meta-analysis. Animal Behaviour, 77(4), 771–783. 10.1016/j.anbehav.2008.12.022

Bierbach, D., Laskowski, K. L., & Wolf, M. (2017). Behavioural individuality in clonal fish arises despite near-identical rearing conditions. Nature Communications, 8(1), 15361. 10.1038/ncomms15361

Biro, P. A. (2013). On the use of rapid assays in personality research: a response to Edwards et al. Animal Behaviour, 86(1), e1–e3. 10.1016/j.anbehav.2013.04.020

Biro, P. A., & Stamps, J. A. (2015). Using repeatability to study physiological and behavioural traits: ignore time-related change at your peril. Animal Behaviour, 105, 223–230. 10.1016/j.anbehav.2015.04.008

Bro-Jørgensen, J., Franks, D. W., & Meise, K. (2019). Linking behaviour to dynamics of populations and communities: application of novel approaches in behavioural ecology to conservation. Philosophical Transactions of the Royal Society B: Biological Sciences, 374(1781), 20190008. 10.1098/rstb.2019.0008

Bruijning, M., Metcalf, C. J. E., Jongejans, E., & Ayroles, J. F. (2020). The evolution of variance control. Trends in Ecology & Evolution, 35(1), 22–33. 10.1016/j.tree.2019.08.005

Bürkner, P. C. (2013). brms: an R package for Bayesian multilevel models using Stan. Journal of Statistical Software, 80(Plummer), 1–28. 10.18637/jss.v080.i01

Cleasby, I. R., Nakagawa, S., & Schielzeth, H. (2015). Quantifying the predictability of behaviour: statistical approaches for the study of between-individual variation in the within-individual variance. Methods in Ecology and Evolution, 6(1), 27–37. 10.1111/2041-210X.12281

de Bivort, B., Buchanan, S., Skutt-Kakaria, K., Gajda, E., Ayroles, J., O’Leary, C., Reimers, P., Akhund-Zade, J., Senft, R., Maloney, R., Ho, S., Werkhoven, Z., & Smith, M. A.-Y. (2022). Precise quantification of behavioral individuality from 80 million decisions across 183,000 flies. Frontiers in Behavioral Neuroscience, 16, 836626. 10.3389/fnbeh.2022.836626

Dingemanse, N. J., & Dochtermann, N. A. (2013). Quantifying individual variation in behaviour: mixed-effect modelling approaches. The Journal of Animal Ecology, 82(1), 39–54. 10.1111/1365-2656.12013

Dingemanse, N. J., Kazem, A. J. N., Réale, D., & Wright, J. (2010). Behavioural reaction norms: animal personality meets individual plasticity. Trends in Ecology & Evolution, 25(2), 81–89. 10.1016/j.tree.2009.07.013

Dingemanse, N. J., & Réale, D. (2005). Natural selection and animal personality. Behaviour, 142(9–10), 1159–1184. 10.1163/156853905774539445

Ehlman, S. M., Scherer, U., Bierbach, D., Francisco, F. A., Laskowski, K. L., Krause, J., & Wolf, M. (2023). Leveraging big data to uncover the eco-evolutionary factors shaping behavioural development. Proceedings of the Royal Society B: Biological Sciences, 290(1992), 20222115. 10.1098/rspb.2022.2115

Ehlman, S. M., Scherer, U., Bierbach, D., Stärk, L., Beese, M., & Wolf, M. (2025). Developmental arcs of plasticity in whole movement repertoires of a clonal fish. iScience, 28(9), 113189. 10.1016/j.isci.2025.113189

Frankenhuis, W. E., & Panchanathan, K. (2011). Individual differences in developmental plasticity may result from stochastic sampling. Perspectives on Psychological Science, 6(4), 336–347. 10.1177/1745691611412602

Freund, J., Brandmaier, A. M., Lewejohann, L., Kirste, I., Kritzler, M., Krüger, A., Sachser, N., Lindenberger, U., & Kempermann, G. (2013). Emergence of individuality in genetically identical mice. Science, 340(6133), 756–759. 10.1126/science.1235294

Freund, J., Brandmaier, A. M., Lewejohann, L., Kirste, I., Kritzler, M., Krüger, A., Sachser, N., Lindenberger, U., & Kempermann, G. (2015). Association between exploratory activity and social individuality in genetically identical mice living in the same enriched environment. Neuroscience, 309, 140–152. 10.1016/j.neuroscience.2015.05.027

Groothuis, T. G. G., & Trillmich, F. (2011). Unfolding personalities: the importance of studying ontogeny. Developmental Psychobiology, 53(6), 641–655. 10.1002/dev.20574

Grunst, A. S., & Grunst, M. L. (2024). Animal personality in multiple stressor environments: the evolutionary ecology of among-individual differences in responses to stressor suites. Proceedings of the Royal Society B: Biological Sciences, 291(2033), 20241620. 10.1098/rspb.2024.1620

Hayes, J. P., & Jenkins, S. H. (1997). Individual variation in mammals. Journal of Mammalogy, 78(2), 274–293. 10.2307/1382882

Honegger, K., & de Bivort, B. (2018). Stochasticity, individuality and behavior. Current Biology, 28(1), R8–R12. 10.1016/j.cub.2017.11.058

Lamatsch, D. K., Schmid, M., & Schartl, M. (2002). A somatic mosaic of the gynogenetic Amazon molly. Journal of Fish Biology, 60(6), 1417–1422. 10.1111/j.1095-8649.2002.tb02436.x

Laskowski, K. L., Bierbach, D., Jolles, J. W., Doran, C., & Wolf, M. (2022). The emergence and development of behavioral individuality in clonal fish. Nature Communications, 13(1), 6419. 10.1038/s41467-022-34113-y

Laskowski, K. L., Chang, C.-C., Sheehy, K., & Aguiñaga, J. (2022). Consistent individual behavioral variation: what do we know and where are we going? Annual Review of Ecology, Evolution, and Systematics, 53(1), 161–182. 10.1146/annurev-ecolsys-102220-011451

MacKinlay, R. D., & Shaw, R. C. (2023). A systematic review of animal personality in conservation science. Conservation Biology, 37(1), e13935. 10.1111/cobi.13935

Mitchell, D. J., Beckmann, C., & Biro, P. A. (2021). Understanding the unexplained: the magnitude and correlates of individual differences in residual variance. Ecology and Evolution, 11(12), 7201–7210. 10.1002/ece3.7603

Mitchell, D. J., Dujon, A. M., Beckmann, C., & Biro, P. A. (2020). Temporal autocorrelation: a neglected factor in the study of behavioral repeatability and plasticity. Behavioral Ecology, 31(1), 222–231. 10.1093/beheco/arz180

Molenaar, P. C. M., Boomsma, D. I., & Dolan, C. V. (1993). A third source of developmental differences. Behavior Genetics, 23(6), 519–524. 10.1007/BF01068142

Mulder, H. A., Gienapp, P., & Visser, M. E. (2016). Genetic variation in variability: phenotypic variability of fledging weight and its evolution in a songbird population. Evolution, 70(9), 2004–2016. 10.1111/evo.13008

Nakagawa, S., & Schielzeth, H. (2010). Repeatability for Gaussian and non-Gaussian data: a practical guide for biologists. Biological Reviews, 85(4), 935–956. 10.1111/j.1469-185X.2010.00141.x

Niemelä, P. T., & Dingemanse, N. J. (2017). Individual versus pseudo-repeatability in behaviour: lessons from translocation experiments in a wild insect. Journal of Animal Ecology, 86(5), 1033–1043. 10.1111/1365-2656.12688

O’Dea, R. E., Noble, D. W. A., & Nakagawa, S. (2021). Unifying individual differences in personality, predictability, and plasticity: a practical guide. Methods in Ecology and Evolution, 2021(January), 1–16. 10.1111/2041-210x.13755

Polverino, G., Cigliano, C., Nakayama, S., & Mehner, T. (2016). Emergence and development of personality over the ontogeny of fish in absence of environmental stress factors. Behavioral Ecology and Sociobiology, 70(12), 2027–2037. 10.1007/s00265-016-2206-z

R Core Team. (2023). R: a language and environment for statistical computing. R Foundation for Statistical Computing. https://www.R-project.org/

Raj, A., & van Oudenaarden, A. (2008). Nature, nurture, or chance: stochastic gene expression and its consequences. Cell, 135(2), 216–226. 10.1016/j.cell.2008.09.050

Réale, D., Reader, S. M., Sol, D., McDougall, P. T., & Dingemanse, N. J. (2007). Integrating animal temperament within ecology and evolution. Biological Reviews, 82(2), 291–318. 10.1111/j.1469-185X.2007.00010.x

Roberts, B. W., & DelVecchio, W. F. (2000). The rank-order consistency of personality traits from childhood to old age: a quantitative review of longitudinal studies. Psychological Bulletin, 126(1), 3–25. 10.1037/0033-2909.126.1.3

Sánchez-Tójar, A., Moiron, M., & Niemelä, P. T. (2022). Terminology use in animal personality research: a self-report questionnaire and a systematic review. Proceedings of the Royal Society B: Biological Sciences, 289(1968), 20212259. 10.1098/rspb.2021.2259

Scherer, U., Ehlman, S. M., Bierbach, D., Krause, J., & Wolf, M. (2023). Reproductive individuality of clonal fish raised in near-identical environments and its link to early-life behavioral individuality. Nature Communications, 14(1), 7652. 10.1038/s41467-023-43069-6

Scherer, U., Laskowski, K. L., Kressler, M. M., Ehlman, S. M., Wolf, M., & Bierbach, D. (2024). Predator exposure early in life shapes behavioral development and individual variation in a clonal fish. Scientific Reports, 14(1), 21668. 10.1038/s41598-024-72550-5

Schlupp, I. (2005). The evolutionary ecology of gynogenesis. Annual Review of Ecology, Evolution, and Systematics, 36, 399–417. 10.1146/annurev.ecolsys.36.102003.152629

Shrout, P. E., & Fleiss, J. L. (1979). Intraclass correlations: uses in assessing rater reliability. Psychological Bulletin, 86(2), 420.

Sih, A., Bell, A. M., Johnson, J. C., & Ziemba, R. E. (2004). Behavioral syndromes: an integrative overview. The Quarterly Review of Biology, 79(3), 241–277. 10.1086/422893

Stamps, J. A., Briffa, M., & Biro, P. A. (2012). Unpredictable animals: individual differences in intraindividual variability (IIV). Animal Behaviour, 83(6), 1325–1334. 10.1016/j.anbehav.2012.02.017

Stamps, J. A., & Groothuis, T. G. G. (2010a). Developmental perspectives on personality: implications for ecological and evolutionary studies of individual differences. Philosophical Transactions of the Royal Society B: Biological Sciences, 365(1560), 4029–4041. 10.1098/rstb.2010.0218

Stamps, J. A., & Groothuis, T. G. G. (2010b). The development of animal personality: relevance, concepts and perspectives. Biological Reviews, 85(2), 301–325. 10.1111/j.1469-185X.2009.00103.x

Stoffel, M. A., Nakagawa, S., & Schielzeth, H. (2017). rptR: repeatability estimation and variance decomposition by generalized linear mixed-effects models. Methods in Ecology and Evolution, 8(11), 1639–1644. 10.1111/2041-210X.12797

Trillmich, F., Müller, T., & Müller, C. (2018). Understanding the evolution of personality requires the study of mechanisms behind the development and life history of personality traits. Biology Letters, 14(2), 20170740. 10.1098/rspb.2017.0740

van Oers, K., & Mueller, J. C. (2010). Evolutionary genomics of animal personality. Philosophical Transactions of the Royal Society B: Biological Sciences, 365(1560), 3991–4000. 10.1098/rstb.2010.0178

Vogt, G., Huber, M., Thiemann, M., van den Boogaart, G., Schmitz, O. J., & Schubart, C. D. (2008). Production of different phenotypes from the same genotype in the same environment by developmental variation. Journal of Experimental Biology, 211(4), 510–523. 10.1242/jeb.008755

Walasek, N., Bruijning, M., Panchanathan, K., & Frankenhuis, W. E. (2026). Noisy information about the environment: a source of individual differences within and across generations. bioRxiv, preprint. 10.64898/2026.05.15.725332

Walasek, N., Jovicic, M., & Guenther, A. (2025). Personality development in non-domesticated house mice: evidence for a nutrition-dependent sensitive period early in life. Proceedings of the Royal Society B: Biological Sciences, 292(2050), 20242689. 10.1098/rspb.2024.2689

Walasek, N., Panchanathan, K., & Frankenhuis, W. E. (2025). The evolution of sensitive periods beyond early ontogeny: bridging theory and data. Functional Ecology, 39(11), 2962–2973. 10.1111/1365-2435.14615

West-Eberhard, M. J. (2003). Developmental plasticity and evolution. Oxford University Press.

Westneat, D. F., Wright, J., & Dingemanse, N. J. (2015). The biology hidden inside residual within individual phenotypic variation. Biological Reviews, 90(3), 729–743. 10.1111/brv.12131

Wolak, M. E., Fairbairn, D. J., & Paulsen, Y. R. (2012). Guidelines for estimating repeatability. Methods in Ecology and Evolution, 3(1), 129–137. 10.1111/j.2041-210X.2011.00125.x

Wolf, M., & Weissing, F. J. (2012). Animal personalities: consequences for ecology and evolution. Trends in Ecology & Evolution, 27(8), 452–461. 10.1016/j.tree.2012.05.001

Wuerz, Y., & Krüger, O. (2015). Personality over ontogeny in zebra finches: long-term repeatable traits but unstable behavioural syndromes. Frontiers in Zoology, 12(1). 10.1186/1742-9994-12-S1-S9

