## supplementary for "The ontogeny of repeatability: measurement choices shape individuality estimates in a clonal fish"

Systematic search for empirical studies on animal personality and repeatability

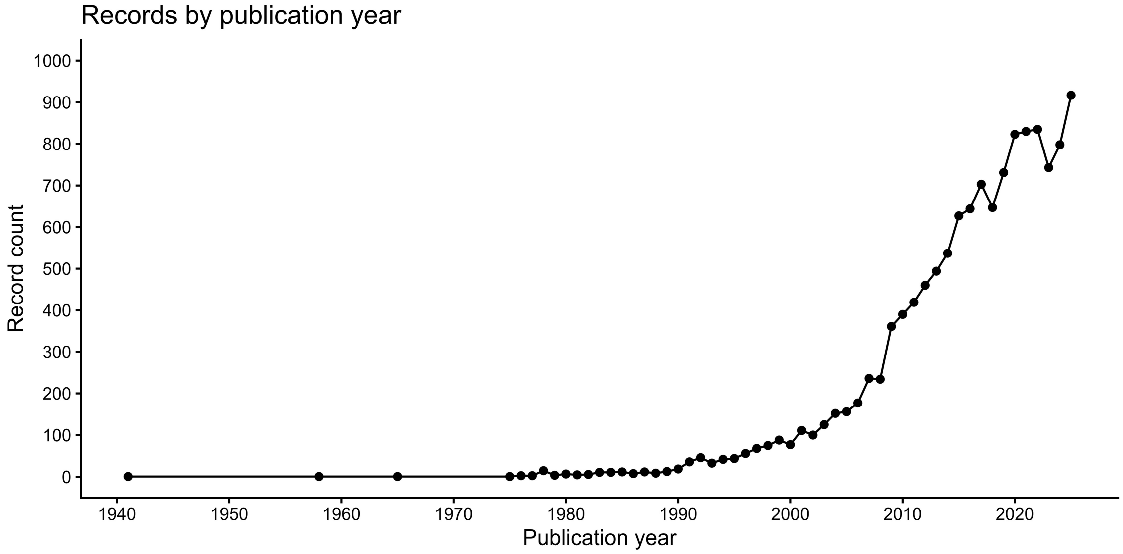


*Figure S1:* Number of articles studying individuality/animal personality across years until the end of 2025. Results have been obtained from a systematic search on Web of Knowledge conducted on 21.08.2026 using the search term: “ALL=(biology, ecology) AND DT=(Article) AND ALL=(animal behavior OR animal personality OR repeatability) AND ALL=(empirical OR study OR experiment OR test)”.

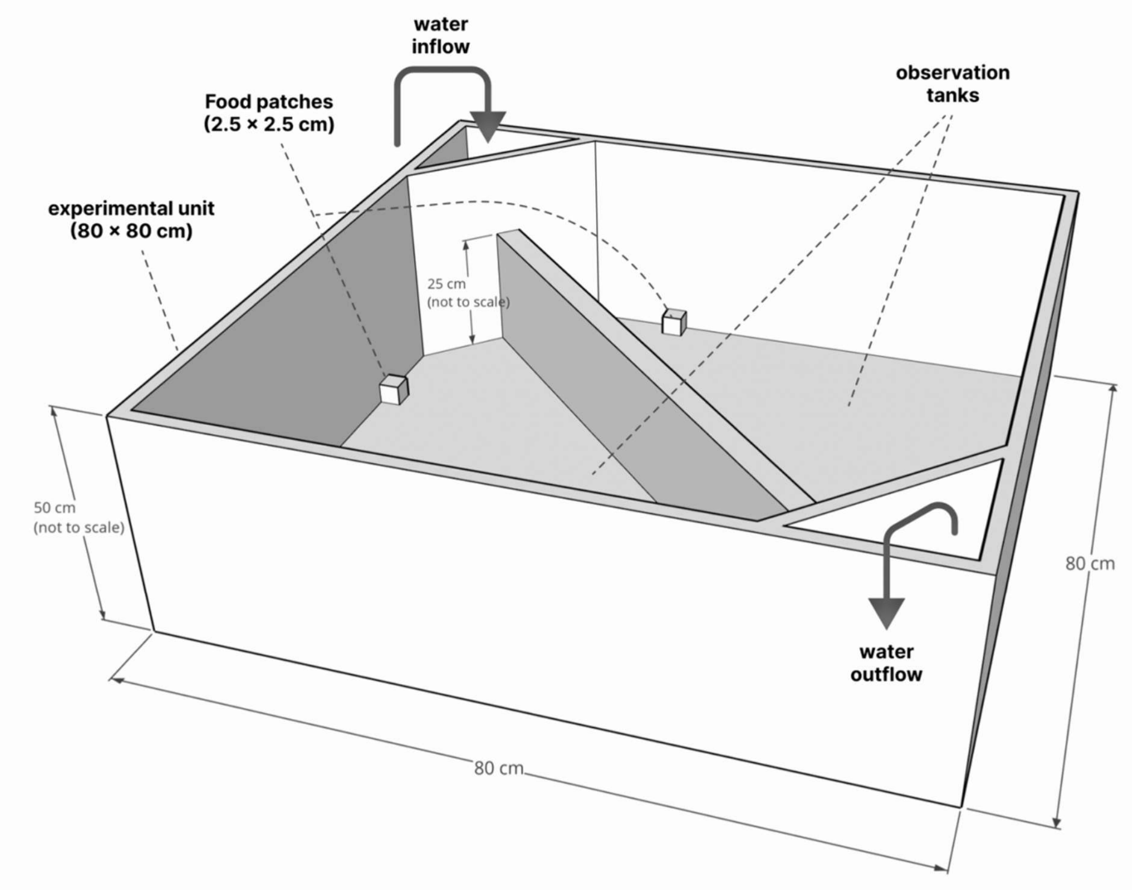


*Figure S2:* Graphical representation of the fish tank. One experimental unit with 2 observation tanks. Water level in the tanks: 7 cm. Food patches were present during the feeding only. Observation tanks were illuminated individually from below with 4 LEDs per tank (each LED is 100cm, 12V, color temperature = 5500 K, light output = approx. 1570 lumen; tanks were manufactured from white polyethylene, which allowed light from underneath to get through). There was no visual contact between observation tanks, but tanks were connected via a flow-through water system (24 observation tanks split into 4 flowthrough systems). Previous work (Scherer et al., 2023) found no significant effect of the tank system (1-4), distance to the external filter unit (0-2m) or centrality of the tank on individual daily activity or feeding.

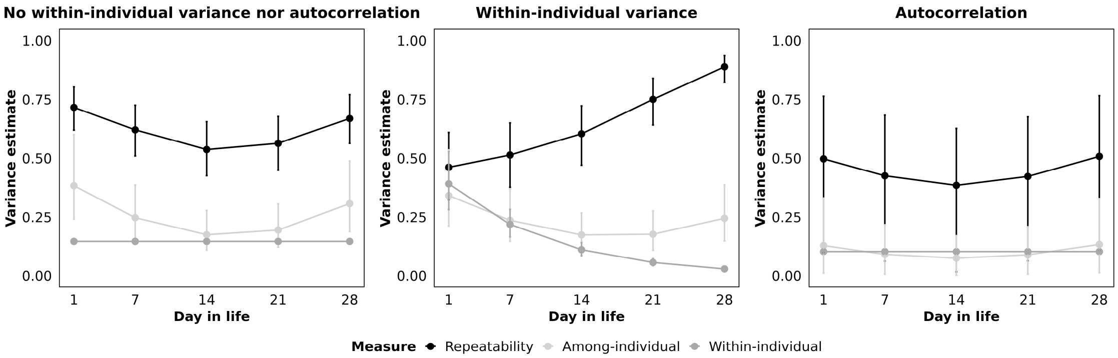


*Figure S3.1:* Sensitivity analysis modeling varying within-individual variation and autocorrelation in daily activity data. Each panel shows variance components (on log-activity scale) and repeatability across ontogeny (vertical axis), estimated from a different mixed effects model. All models included activity on the log-scale as the outcome variable and centered age in days, standard length (cm), and maternal identity as fixed effects. Additionally, we allowed individual-specific trajectories over time by including random intercepts for individuals and random slopes for age. Additionally, models either included varying within-individual variance or autocorrelation.

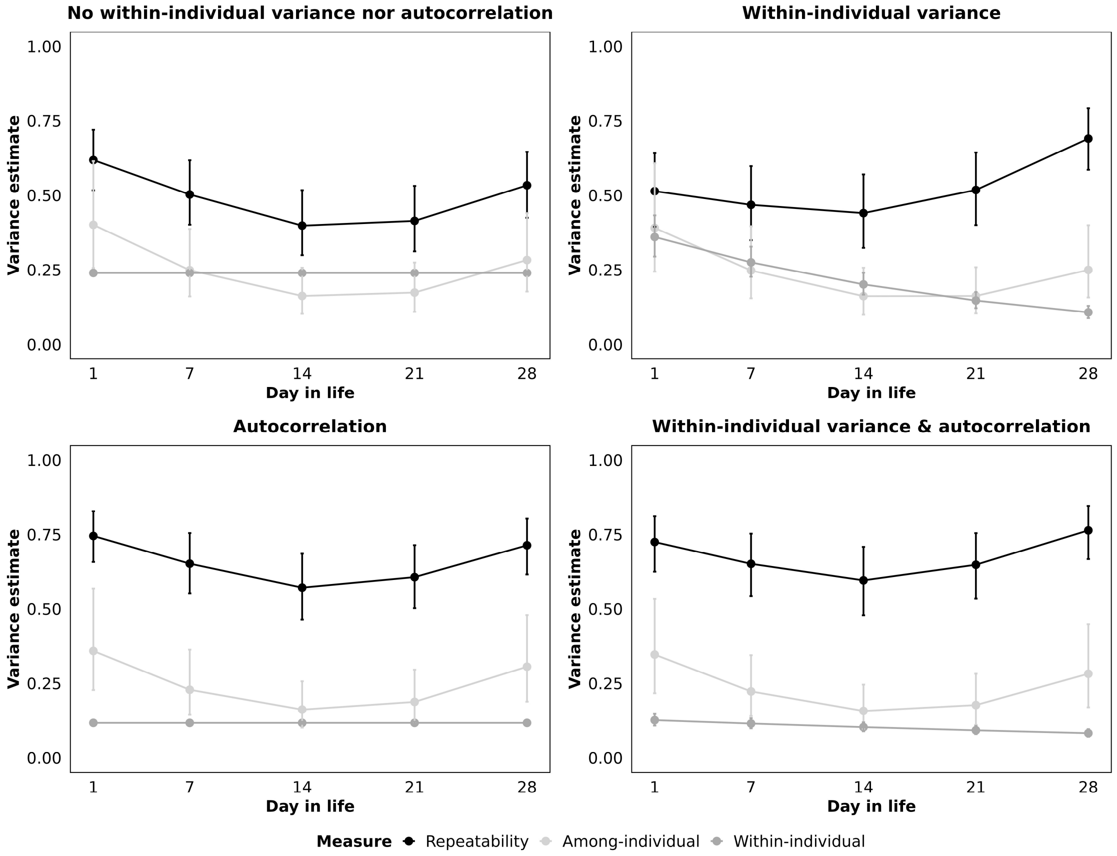


*Figure S3.2:* Sensitivity analysis modeling varying within-individual variation and autocorrelation in hourly activity data. Each panel shows variance components (on log-activity scale) and repeatability across ontogeny (vertical axis), estimated from a different mixed effects model. All models included activity on the log-scale as the outcome variable and centered age in days, standard length (cm), and maternal identity as fixed effects. Additionally, we allowed individual-specific trajectories over time by including random intercepts for individuals and random slopes for age. Additionally, models either included varying within-individual variance, autocorrelation, or both.

<u>Additional results for different sampling rates</u>

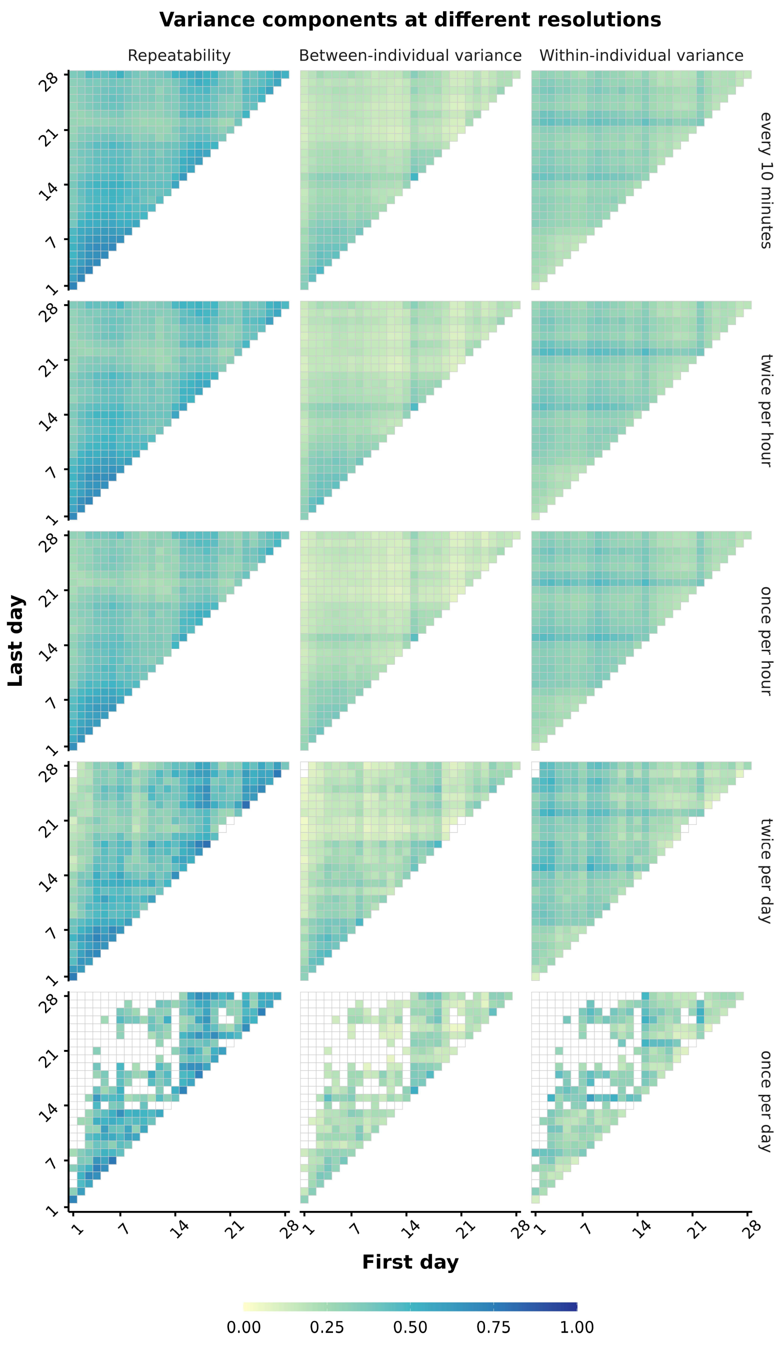


*Figure S4:* Repeatability and variance components across sampling rates. The raw activity observations (recorded at 5 frames per second) were first aggregated across 10-minute observation intervals, resulting in one activity data point per every 10 minutes, per fish. Rows indicate the sampling rate of these 10-minute observation intervals: continuously every 10 minutes, twice per hour, once per hour, twice per day, and once per day. For example, the *twice per hour* sampling rate means that the model included aggregated activity data from two 10-minute observation intervals per hour, per individual. Columns show, from left to right, heatmaps illustrating repeatability ($R$), among-individual variance, and within-individual variance estimates; variance components are reported on the log-activity scale. Each cell within a heatmap represents a significant estimate ($\alpha < .05$) from a separate model fitted to data from the pair of days indicated by the axes. The horizontal axis gives the first day and the vertical axis gives the second day; only unique day pairs are shown. Cells on the diagonal therefore represent models fitted to observations from a single day, whereas off-diagonal cells represent models fitted to observations from two different days. Cell color denotes the estimated value. At the daily aggregation level, estimates from same-day models cannot be obtained because each individual contributes only one observation per day at this aggregation level. Empty cells elsewhere indicate estimates that did not reach the statistical significance threshold. Repeatability is bounded between 0 and 1. Variance components are not inherently bounded, but all estimated values in this dataset were below 1; therefore, a common color scale is used across panels.

<u>Results including all intermediate days</u>

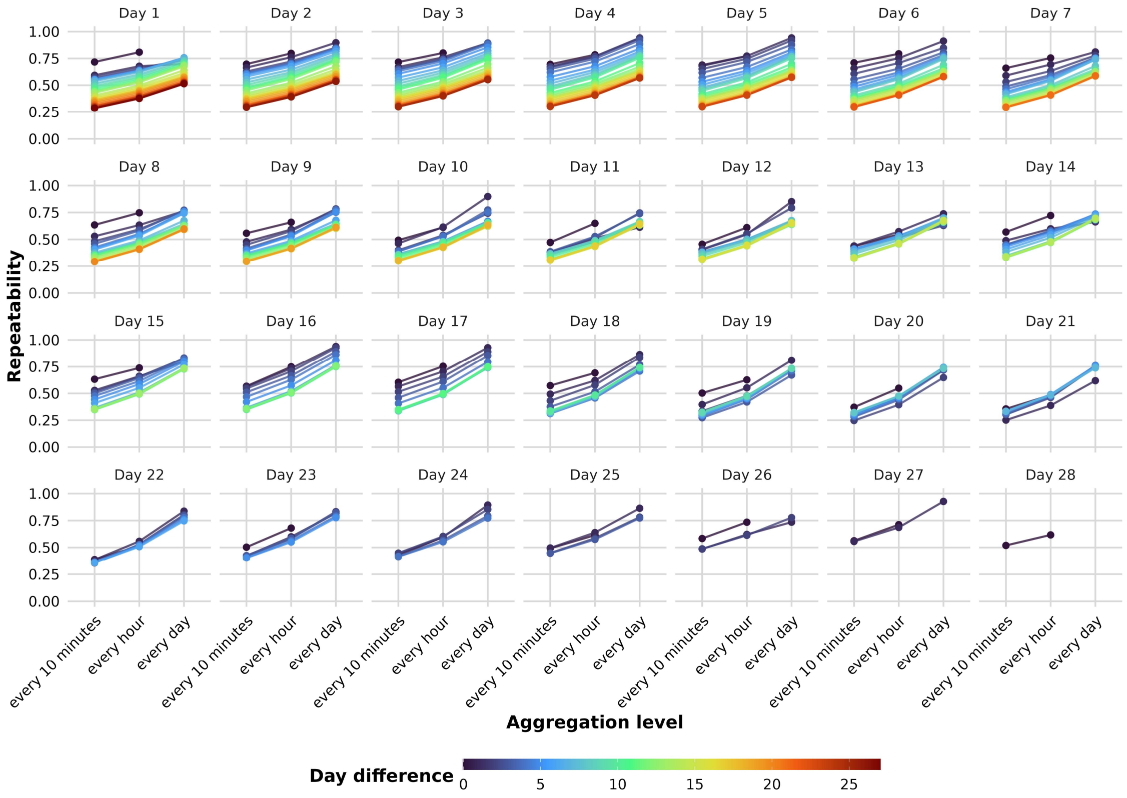

*Figure S5:* Repeatability across aggregation levels computed including intermediate days. Each panel represents a focal day in the fish's life. The vertical axis shows statistically significant ($\alpha < .05$) repeatability estimates. Each filled circle corresponds to one model fitted to data from that focal day and, except where the day difference was zero, some subsequent day (separated from a focal day by the indicated day difference), including all of the intermediate days. Colors of circles and connecting lines indicate the number of days separating the two measurement days (day difference; 0–27 days). The horizontal axis indicates the temporal aggregation level applied to the activity data: every minute, every 10 minutes, every hour, or every day. For example, at the *every minute* level, raw activity observations (recorded at 5 frames per second) were averaged within each minute, yielding one observation per individual per minute. A day difference of zero denotes estimates based on data from the focal day alone. Repeatability could not be estimated at the daily aggregation level for these models because this aggregation yields only one observation per individual per day. Other missing estimates indicate that the repeatability estimate did not reach the statistical significance threshold.

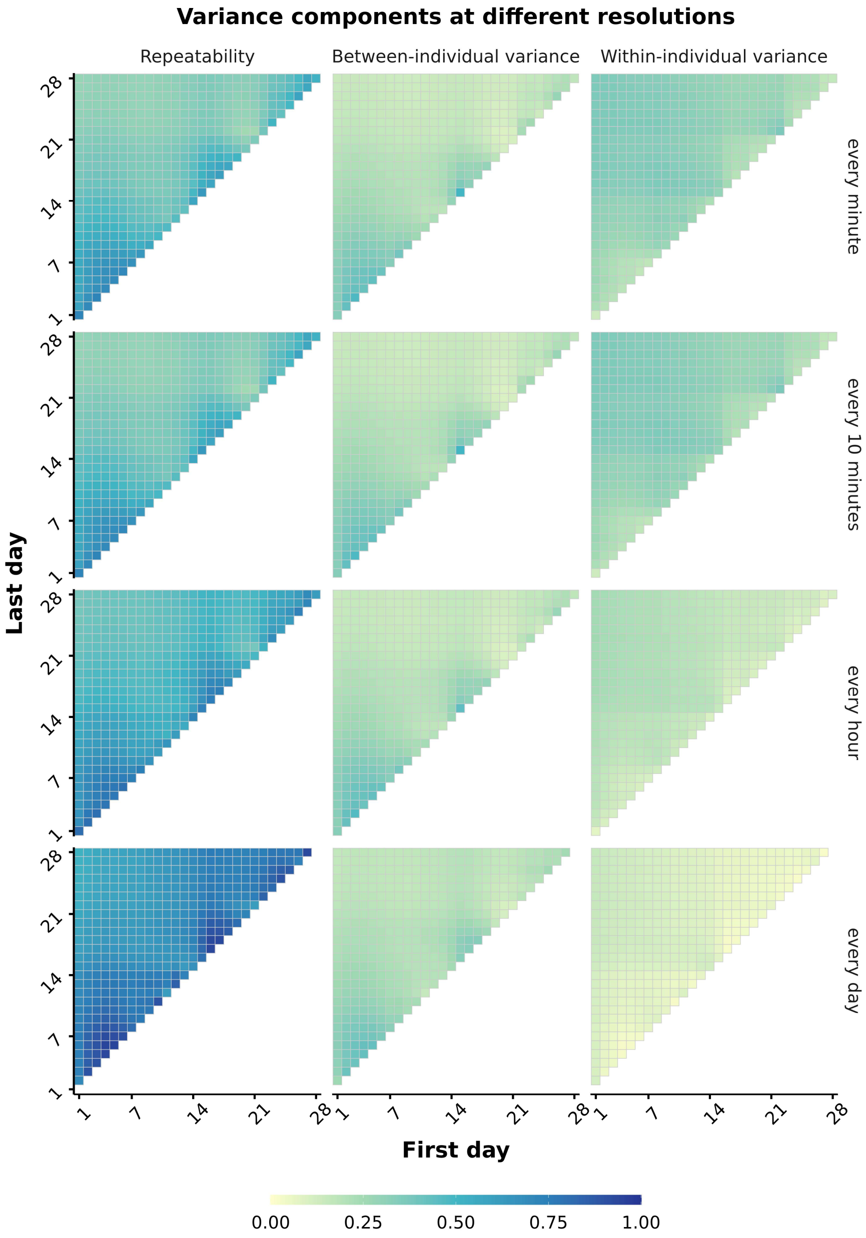


*Figure S6:* Repeatability and variance components across aggregation levels computed including intermediate days. Rows represent the temporal aggregation level applied to the activity data: every minute, every 10 minutes, every hour, and every day. For example, at the every minute level, raw activity observations (recorded at 5 frames per second) were averaged within each minute, yielding one observation per individual per minute. Columns show, from left to right, heatmaps illustrating repeatability ($R$), among-individual variance, and within-individual variance estimates; variance components are reported on the log-activity scale. Each cell within a heatmap represents a significant estimate ($\alpha < .05$) from a separate model fitted to data from the pair of days indicated by the axes, including all intermediate days. The horizontal axis gives the first day and the vertical axis gives the second day; only unique day pairs are shown. Cells on the diagonal therefore represent models fitted to observations from a single day, whereas off-diagonal cells represent models fitted to observations from two different days. Cell color denotes the estimated value. At the daily aggregation level, estimates from same-day models cannot be obtained because each individual contributes only one observation per day at this aggregation level. Empty cells elsewhere indicate estimates that did not reach the statistical significance threshold. Repeatability is bounded between 0 and 1. Variance components are not inherently bounded, but all estimated values in this dataset were below 1; therefore, a common color scale is used across panels.

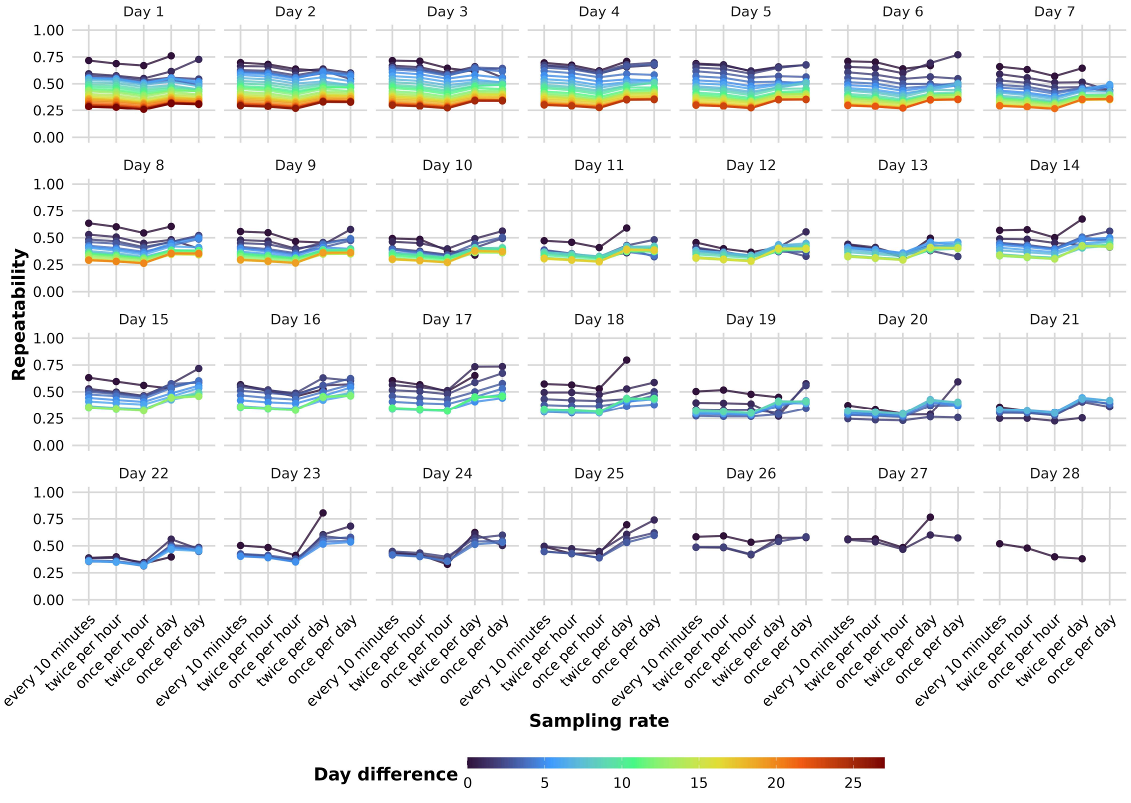


*Figure S7:* Repeatability across sampling rates computed including intermediate days. Each panel represents a focal day in the fish's life. The vertical axis shows statistically significant ($\alpha < .05$) repeatability estimates. Each filled circle corresponds to one model fitted to data from that focal day and, except where the day difference was zero, some subsequent day (separated from a focal day by the indicated day difference), including intermediate days.

Colors of circles and connecting lines indicate the number of days separating the two measurement days (day difference; 0–27 days). The raw activity observations (recorded at 5 frames per second) were first aggregated across 10-minute observation intervals, resulting in one activity data point per every 10 minutes, per fish. The horizontal axis indicates the sampling rate of these 10-minute observation intervals: continuously every 10 minutes, twice per hour, once per hour, twice per day, and once per day. For example, the *twice per hour* sampling rate means that the model included aggregated activity data from two 10-minute observation intervals per hour, per individual. A day difference of zero denotes estimates based on data from the focal day alone. Repeatability could not be estimated at the once per day sampling rate for these models because this rate yields only one observation per individual, per day. Other missing estimates indicate that the repeatability estimate did not reach the statistical significance threshold.

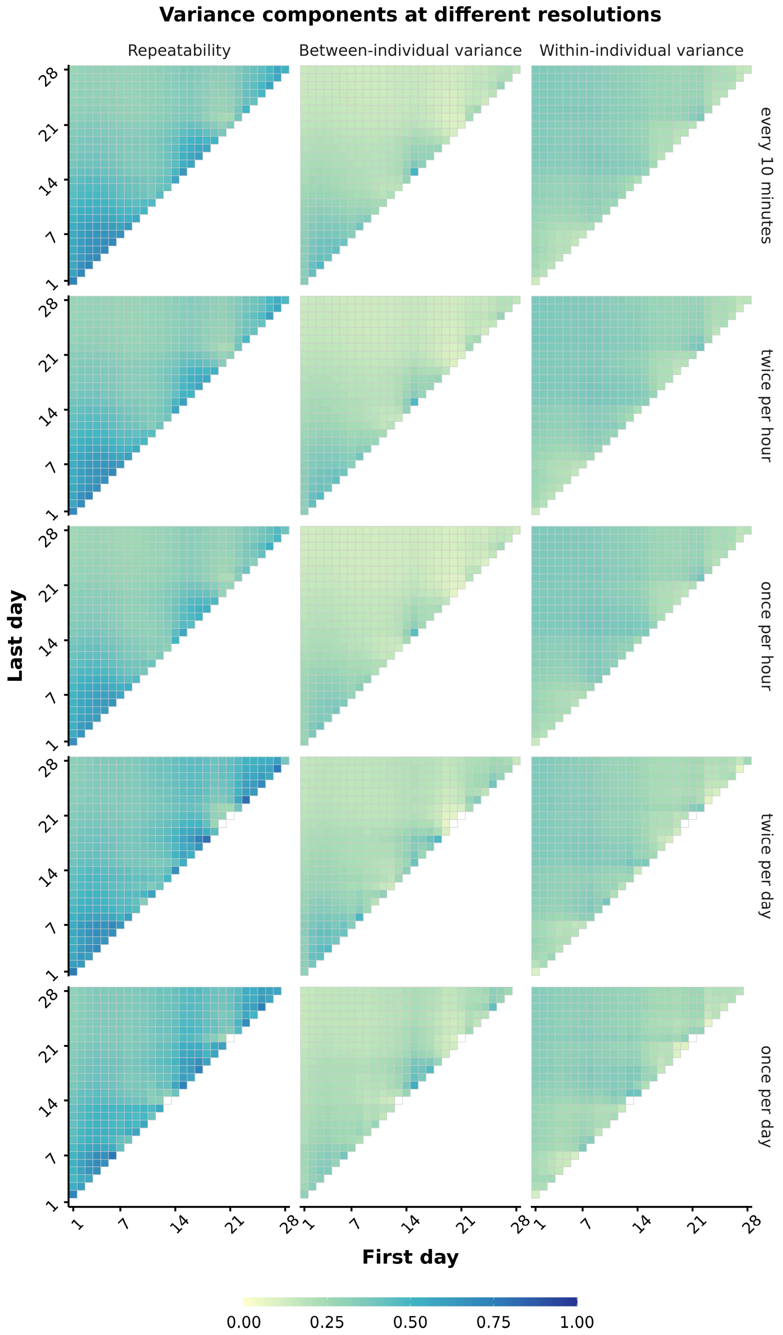


*Figure S8:* Repeatability and variance components across sampling rates computed including intermediate days. The raw activity observations (recorded at 5 frames per second) were first aggregated across 10-minute observation intervals, resulting in one activity data point per every 10 minutes, per fish. Rows indicate the sampling rate of these 10-minute observation intervals: continuously every 10 minutes, twice per hour, once per hour, twice per day, and once per day. For example, the *twice per hour* sampling rate means that the model included aggregated activity data from two 10-minute observation intervals per hour, per individual.

Columns show, from left to right, heatmaps illustrating repeatability ($R$), among-individual variance, and within-individual variance estimates; variance components are reported on the log-activity scale. Each cell within a heatmap represents a significant estimate ($\alpha < .05$) from a separate model fitted to data from the pair of days indicated by the axes, including intermediate days. The horizontal axis gives the first day and the vertical axis gives the second day; only unique day pairs are shown. Cells on the diagonal therefore represent models fitted to observations from a single day, whereas off-diagonal cells represent models fitted to observations from two different days. Cell color denotes the estimated value. At the daily aggregation level, estimates from same-day models cannot be obtained because each individual contributes only one observation per day at this aggregation level. Empty cells elsewhere indicate estimates that did not reach the statistical significance threshold. Repeatability is bounded between 0 and 1. Variance components are not inherently bounded, but all estimated values in this dataset were below 1; therefore, a common color scale is used across panels.

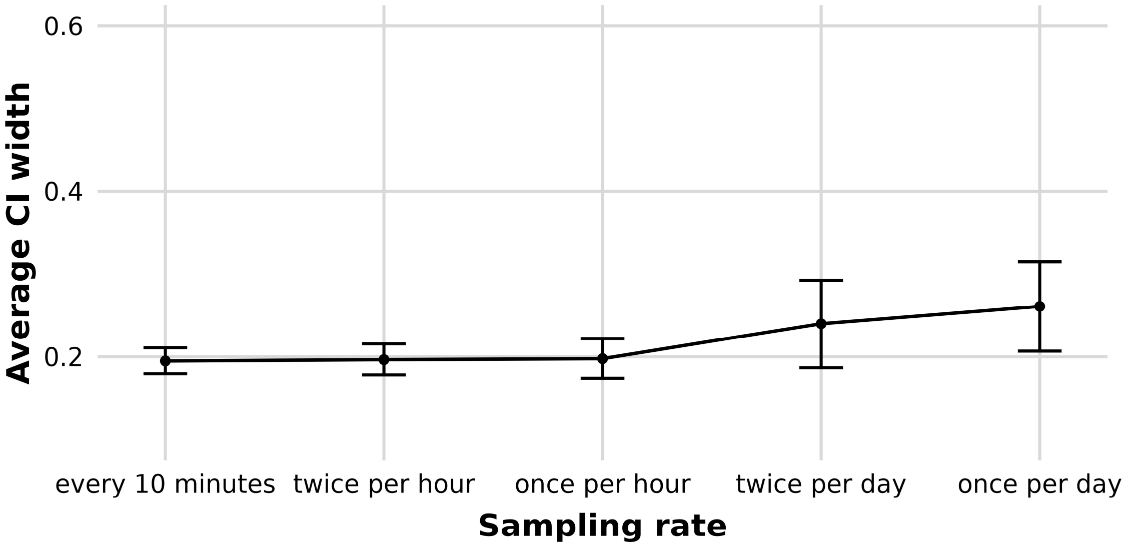


*Figure S9:* Confidence intervals around repeatability estimates across sampling rates computed including intermediate days. 95% confidence intervals are based on models fitted to activity data resulting from different sampling rates of 10-minute observation intervals. The horizontal axis shows these different sampling rates. The vertical axis shows the average width of the confidence intervals across all models computed from data at a specific sampling rate. The error bar indicates the standard deviation around the average width.

Variance decomposition first and last hour of the day measurement times

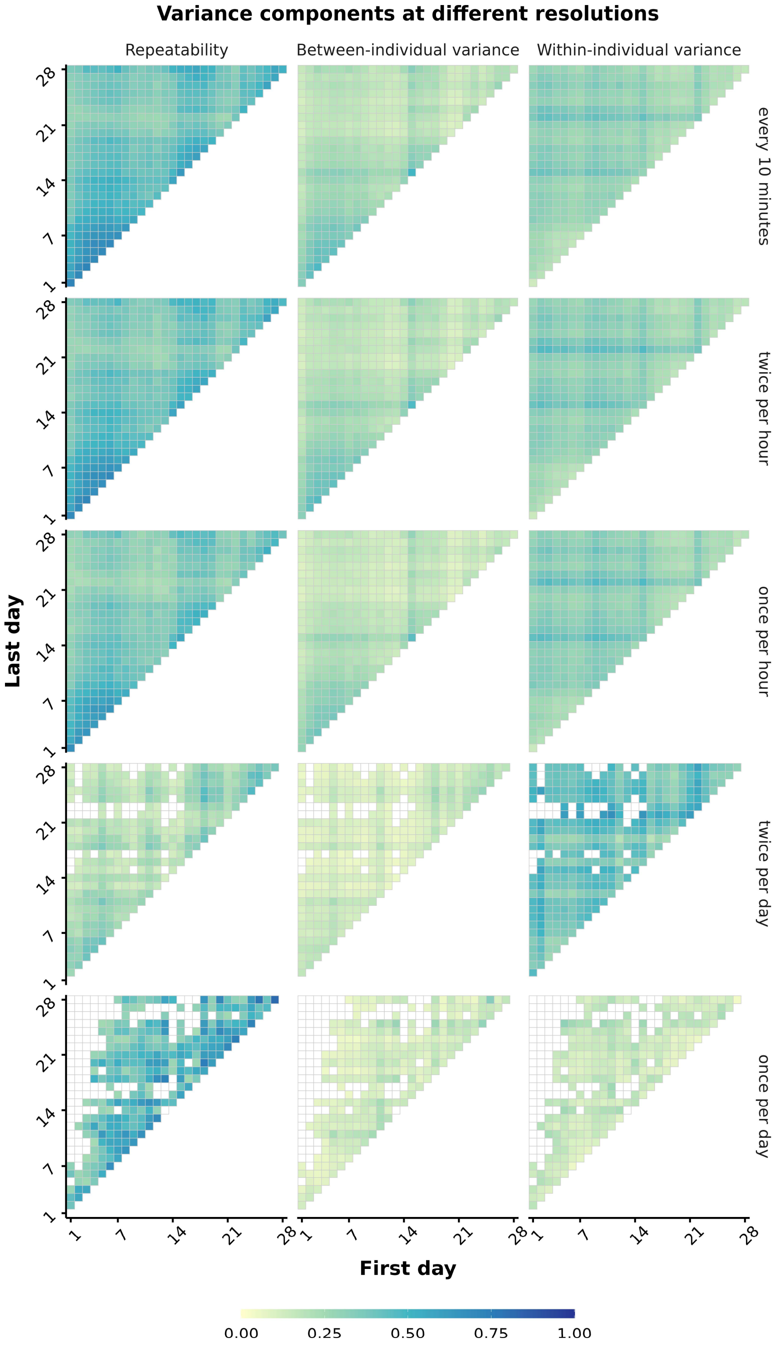


*Figure S10:* Repeatability and variance components across sampling rates and measurement times. The raw activity observations (recorded at 5 frames per second) were first aggregated across 10-minute observation intervals, resulting in one activity data point per every 10 minutes, per fish. Rows indicate the sampling rate of these 10-minute observation intervals:

twice per day using the first and last 10 minutes of each day, twice per day using the second and second-to-last 10 minutes of each day, twice per day using the third and third-to-last 10 minutes of each day, once per day using the first 10 minutes of each day, once per day using the second 10 minutes of each day, and once per day using the third 10 minutes of each day. Columns show, from left to right, heatmaps illustrating repeatability ($R$), among-individual variance, and within-individual variance estimates; variance components are reported on the log-activity scale. Each cell within a heatmap represents a significant estimate ($\alpha < .05$) from a separate model fitted to data from the pair of days indicated by the axes. The horizontal axis gives the first day and the vertical axis gives the second day; only unique day pairs are shown. Cells on the diagonal therefore represent models fitted to observations from a single day, whereas off-diagonal cells represent models fitted to observations from two different days. Cell color denotes the estimated value. At the daily aggregation level, estimates from same-day models cannot be obtained because each individual contributes only one observation per day at this aggregation level. Empty cells elsewhere indicate estimates that did not reach the statistical significance threshold. Repeatability is bounded between 0 and 1. Variance components are not inherently bounded, but all estimated values in this dataset were below 1; therefore, a common color scale is used across panels.

### Results for different frame rates

#### *1 frame per second*

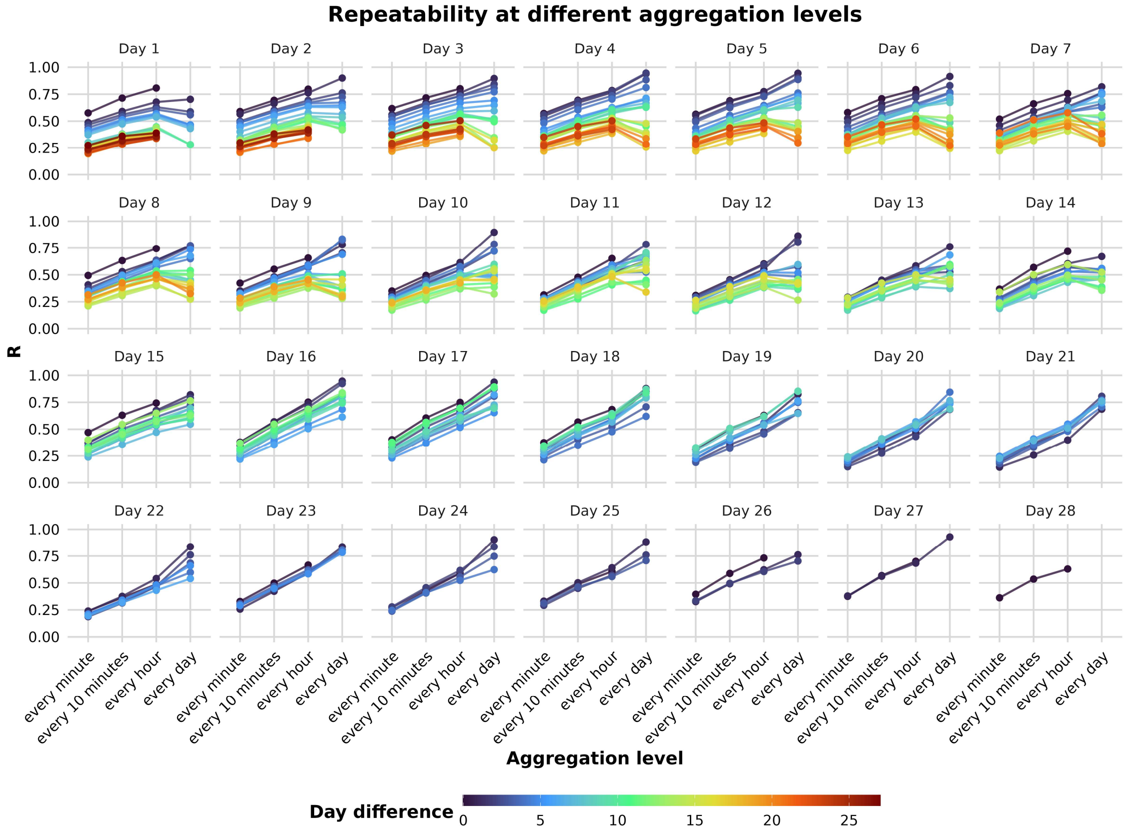


*Figure S11:* Repeatability across aggregation levels computed from activity data tracked at 1 frame per second. Each panel represents a focal day in the fish's life. The vertical axis shows statistically significant ($\alpha < .05$) repeatability estimates. Each filled circle corresponds to one model fitted to data from that focal day and, except where the day difference was zero, some subsequent day (separated from a focal day by the indicated day difference). Colors of circles and connecting lines indicate the number of days separating the two measurement days (day difference; 0–27 days). The horizontal axis indicates the temporal aggregation level applied to the activity data: every minute, every 10 minutes, every hour, or every day. For example, at the *every minute* level, raw activity observations (recorded at 5 frames per second) were averaged within each minute, yielding one observation per individual per minute. A day difference of zero denotes estimates based on data from the focal day alone. Repeatability could not be estimated at the daily aggregation level for these models because this aggregation yields only one observation per individual per day. Other missing estimates indicate that the repeatability estimate did not reach the statistical significance threshold.

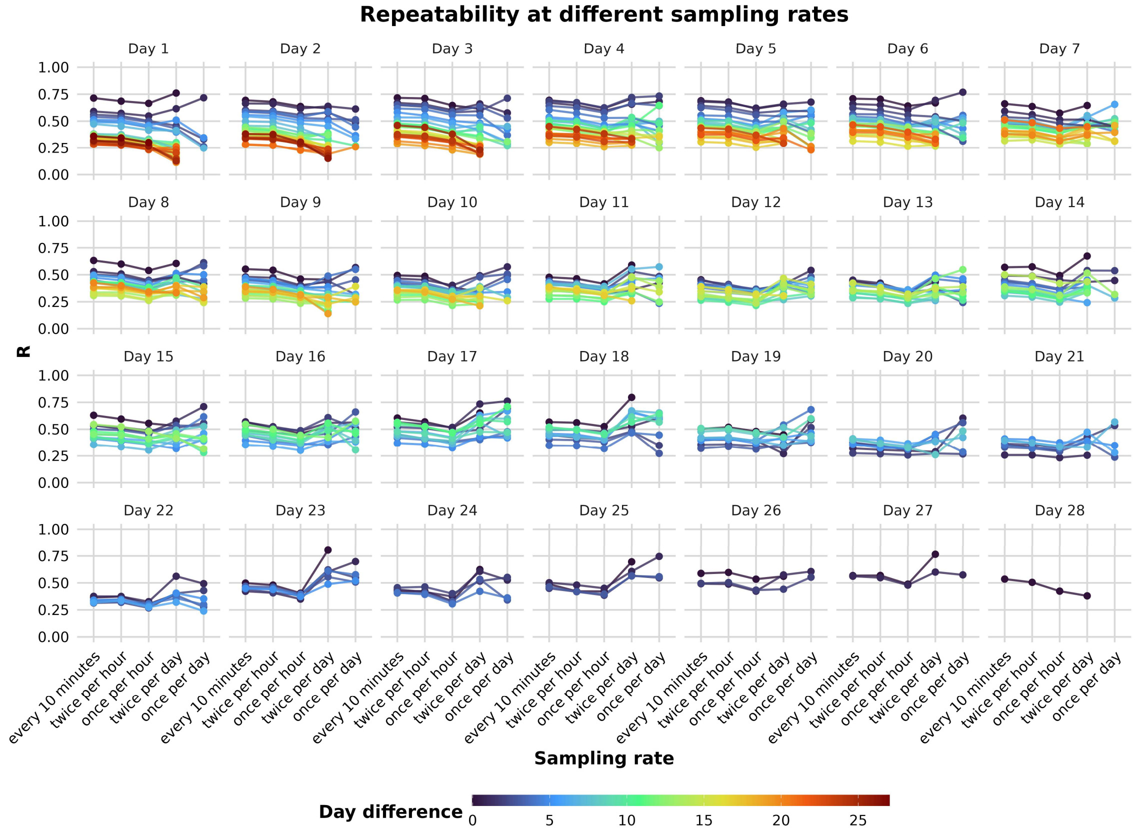


*Figure S12:* Repeatability across sampling rates computed from activity data tracked a 1 frame per second. Each panel represents a focal day in the fish's life. The vertical axis shows statistically significant ($\alpha < .05$) repeatability estimates. Each filled circle corresponds to one model fitted to data from that focal day and, except where the day difference was zero, some subsequent day (separated from a focal day by the indicated day difference). Colors of circles and connecting lines indicate the number of days separating the two measurement days (day difference; 0–27 days). The raw activity observations (recorded at 5 frames per second) were first aggregated across 10-minute observation intervals, resulting in one activity data point per every 10 minutes, per fish. The horizontal axis indicates the sampling rate of these 10-minute observation intervals: continuously every 10 minutes, twice per hour, once per hour, twice per day, and once per day. For example, the *twice per hour* sampling rate means that the model included aggregated activity data from two 10-minute observation intervals per hour, per individual. A day difference of zero denotes estimates based on data from the focal day alone. Repeatability could not be estimated at the once per day sampling rate for these models because this rate yields only one observation per individual, per day. Other missing estimates indicate that the repeatability estimate did not reach the statistical significance threshold.

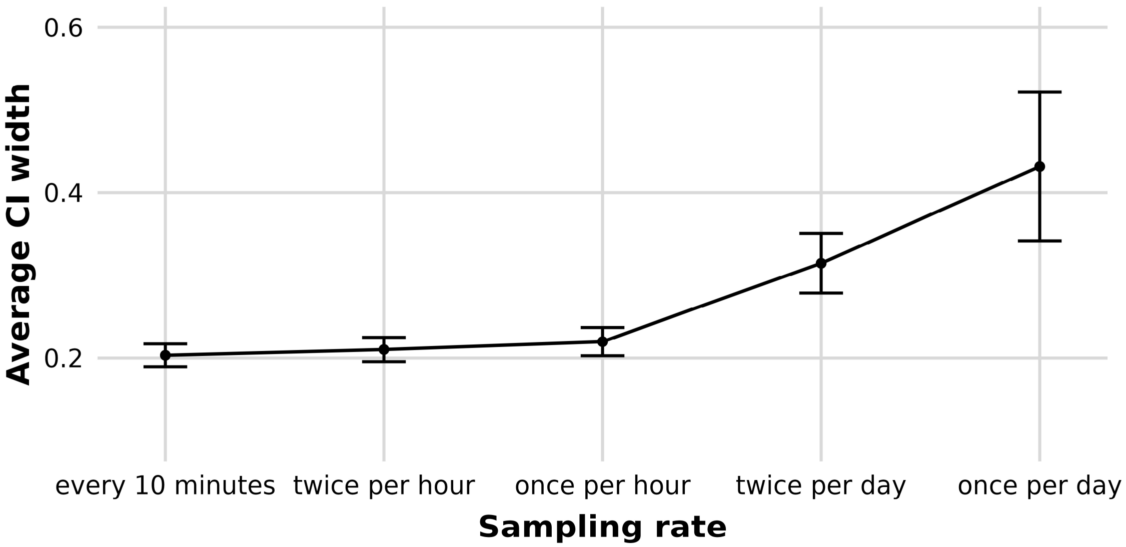


*Figure S13:* Confidence intervals around repeatability estimates across sampling rates computed from activity tracked at 1 frame per second. 95% confidence intervals are based on models fitted to activity data resulting from different sampling rates of 10-minute observation intervals. The horizontal axis shows these different sampling rates. The vertical axis shows the average width of the confidence intervals across all models computed from data at a specific sampling rate. The error bar indicates the standard deviation around the average width.

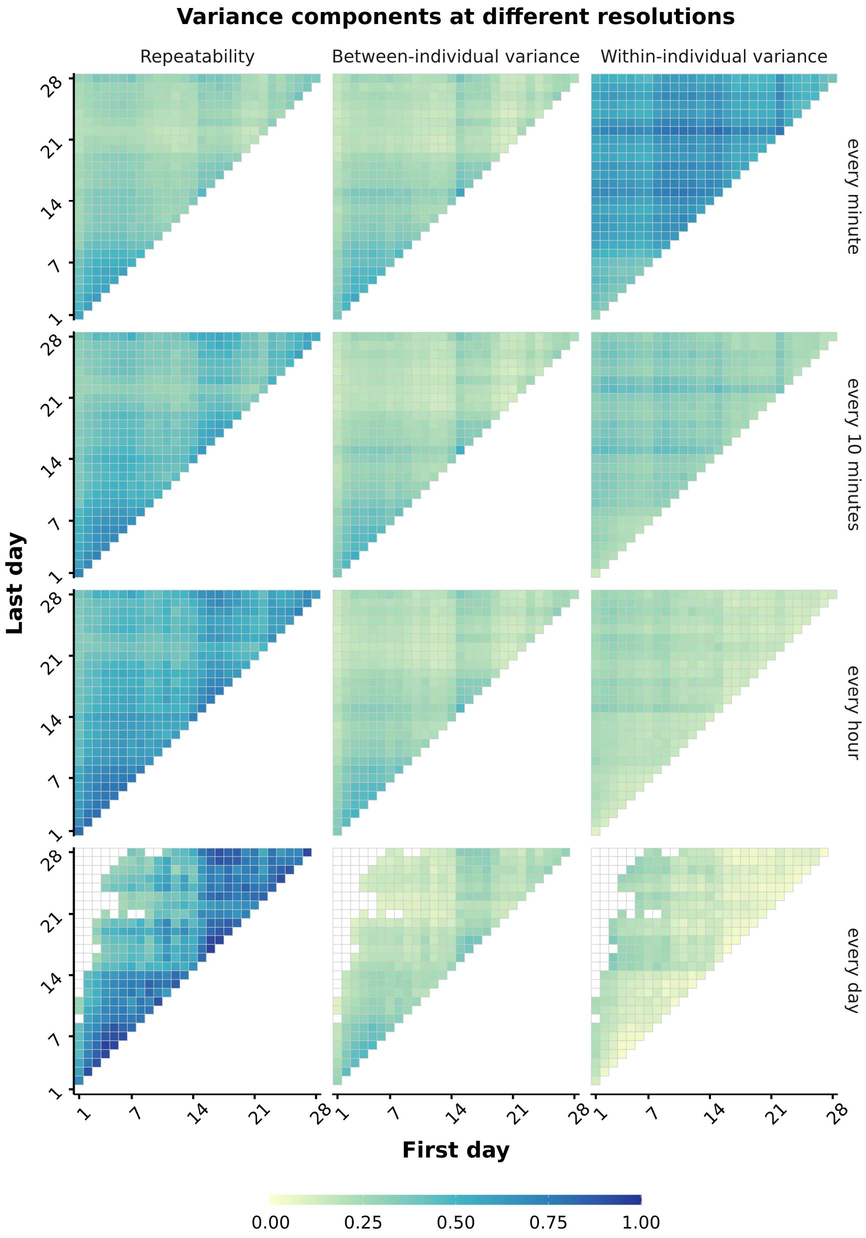


*Figure S14:* Repeatability and variance components across aggregation levels computed from activity data tracked at 1 frame per second. Rows represent the temporal aggregation level applied to the activity data: every minute, every 10 minutes, every hour, and every day. For example, at the every minute level, raw activity observations (recorded at 5 frames per second) were averaged within each minute, yielding one observation per individual per minute. Columns show, from left to right, heatmaps illustrating repeatability ($R$), among-individual variance, and within-individual variance estimates; variance components are reported on the log-activity scale. Each cell within a heatmap represents a significant estimate ($\alpha < .05$) from a separate model fitted to data from the pair of days indicated by the axes. The horizontal axis gives the first day and the vertical axis gives the second day; only unique day pairs are shown. Cells on the diagonal therefore represent models fitted to observations from a single day, whereas off-diagonal cells represent models fitted to observations from two different days. Cell color denotes the estimated value. At the daily aggregation level, estimates from same-day models cannot be obtained because each individual contributes only one observation per day at this aggregation level. Empty cells elsewhere indicate estimates that did not reach the statistical significance threshold. Repeatability is bounded between 0 and 1. Variance components are not inherently bounded, but all estimated values in this dataset were below 1; therefore, a common color scale is used across panels.

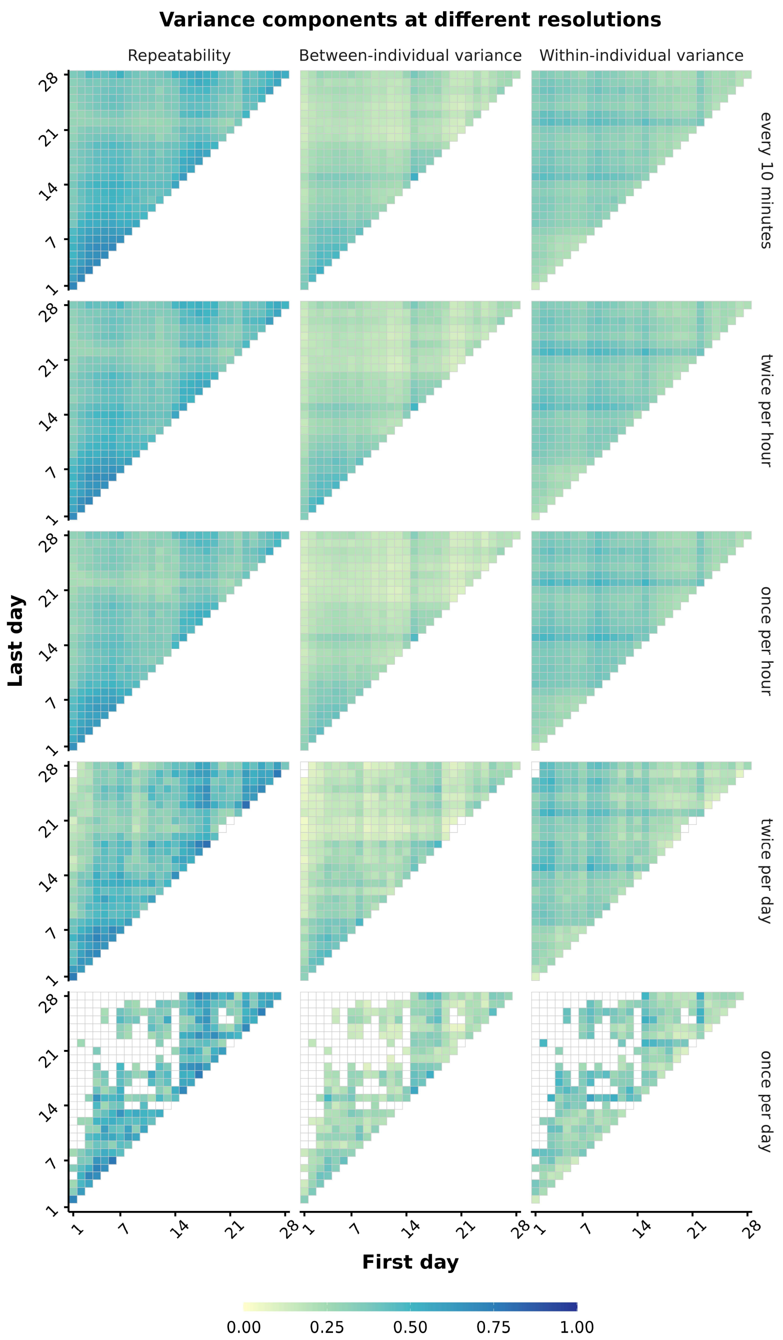


*Figure S15:* Repeatability and variance components across sampling rates computed from activity data tracked at 1 frame per second. The raw activity observations (recorded at 5 frames per second) were first aggregated across 10-minute observation intervals, resulting in one activity data point per every 10 minutes, per fish. Rows indicate the sampling rate of these 10-minute observation intervals: continuously every 10 minutes, twice per hour, once per hour, twice per day, and once per day. For example, the *twice per hour* sampling rate means that the model included aggregated activity data from two 10-minute observation intervals per hour, per individual. Columns show, from left to right, heatmaps illustrating repeatability ($R$), among-individual variance, and within-individual variance estimates; variance components are reported on the log-activity scale. Each cell within a heatmap represents a significant estimate ($\alpha < .05$) from a separate model fitted to data from the pair of days indicated by the axes. The horizontal axis gives the first day and the vertical axis gives the second day; only unique day pairs are shown. Cells on the diagonal therefore represent models fitted to observations from a single day, whereas off-diagonal cells represent models fitted to observations from two different days. Cell color denotes the estimated value. At the daily aggregation level, estimates from same-day models cannot be obtained because each individual contributes only one observation per day at this aggregation level. Empty cells elsewhere indicate estimates that did not reach the statistical significance threshold. Repeatability is bounded between 0 and 1. Variance components are not inherently bounded, but all estimated values in this dataset were below 1; therefore, a common color scale is used across panels.

*1 frame per minute*

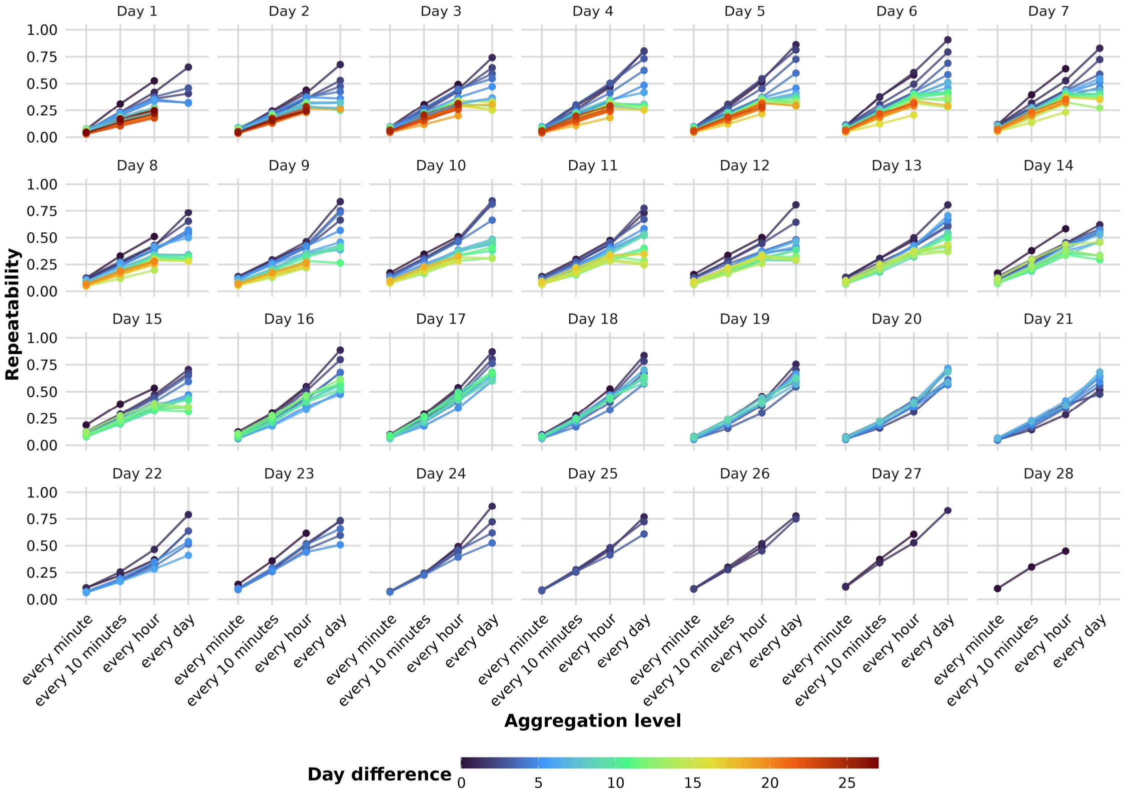

*Figure S16:* Repeatability across aggregation levels computed from activity data tracked at 1 frame per minute. Each panel represents a focal day in the fish's life. The vertical axis shows statistically significant ($\alpha < .05$) repeatability estimates. Each filled circle corresponds to one model fitted to data from that focal day and, except where the day difference was zero, some subsequent day (separated from a focal day by the indicated day difference). Colors of circles and connecting lines indicate the number of days separating the two measurement days (day difference; 0–27 days). The horizontal axis indicates the temporal aggregation level applied to the activity data: every minute, every 10 minutes, every hour, or every day. For example, at the *every minute* level, raw activity observations (recorded at 5 frames per second) were averaged within each minute, yielding one observation per individual per minute. A day difference of zero denotes estimates based on data from the focal day alone. Repeatability could not be estimated at the daily aggregation level for these models because this aggregation yields only one observation per individual per day. Other missing estimates indicate that the repeatability estimate did not reach the statistical significance threshold.

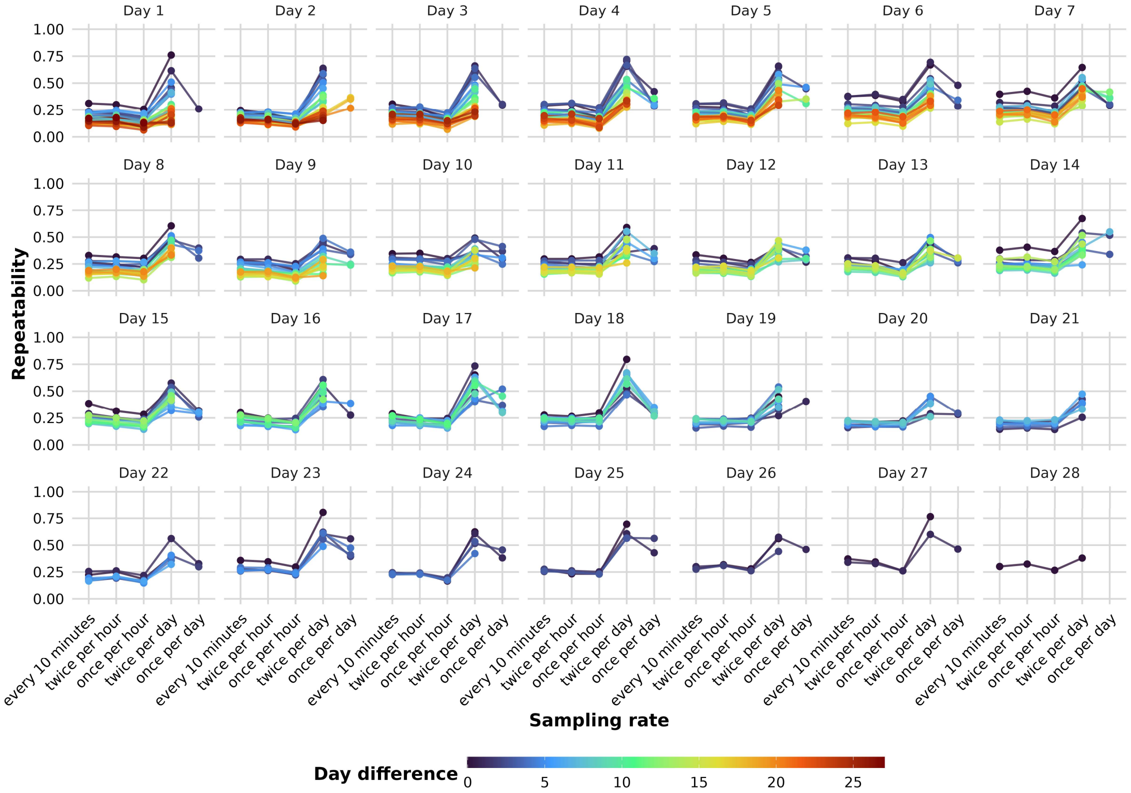

*Figure S17:* Repeatability across sampling rates computed from activity data tracked a 1 frame per minute. Each panel represents a focal day in the fish's life. The vertical axis shows statistically significant ($\alpha < .05$) repeatability estimates. Each filled circle corresponds to one model fitted to data from that focal day and, except where the day difference was zero, some subsequent day (separated from a focal day by the indicated day difference). Colors of circles and connecting lines indicate the number of days separating the two measurement days (day difference; 0–27 days). The raw activity observations (recorded at 5 frames per second) were first aggregated across 10-minute observation intervals, resulting in one activity data point per every 10 minutes, per fish. The horizontal axis indicates the sampling rate of these 10-minute observation intervals: continuously every 10 minutes, twice per hour, once per hour, twice per day, and once per day. For example, the *twice per hour* sampling rate means that the model included aggregated activity data from two 10-minute observation intervals per hour, per individual. A day difference of zero denotes estimates based on data from the focal day alone. Repeatability could not be estimated at the once per day sampling rate for these models because this rate yields only one observation per individual, per day. Other missing estimates indicate that the repeatability estimate did not reach the statistical significance threshold.

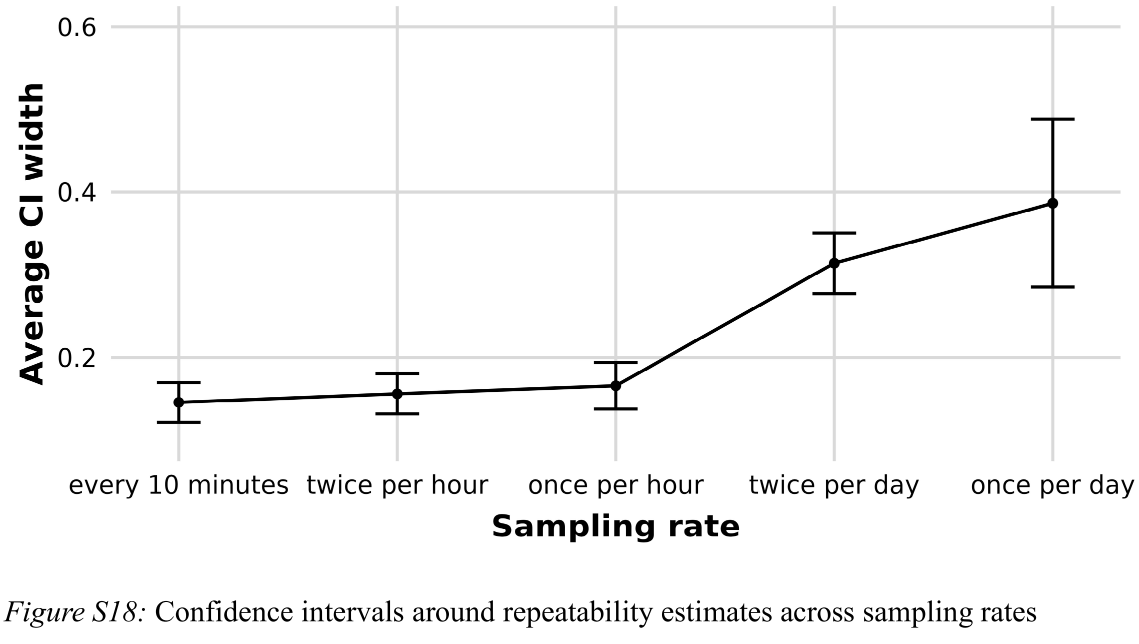


*Figure S18:* Confidence intervals around repeatability estimates across sampling rates computed from activity tracked at 1 frame per minute. 95% confidence intervals are based on models fitted to activity data resulting from different sampling rates of 10-minute observation intervals. The horizontal axis shows these different sampling rates. The vertical axis shows the average width of the confidence intervals across all models computed from data at a specific sampling rate. The error bar indicates the standard deviation around the average width.

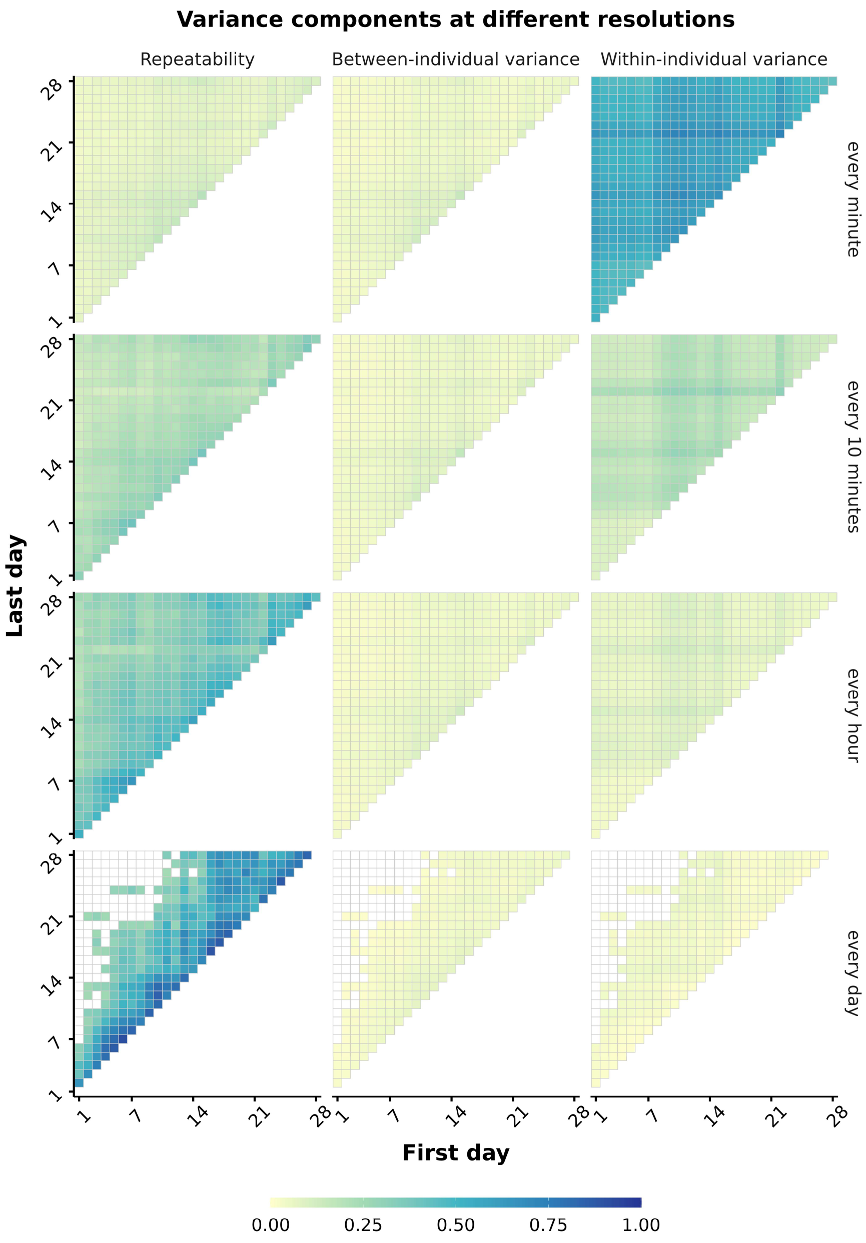


*Figure S19:* Repeatability and variance components across aggregation levels computed from activity data tracked at 1 frame per minute. Rows represent the temporal aggregation level applied to the activity data: every minute, every 10 minutes, every hour, and every day. For example, at the every minute level, raw activity observations (recorded at 5 frames per second) were averaged within each minute, yielding one observation per individual per minute. Columns show, from left to right, heatmaps illustrating repeatability ($R$), among-individual variance, and within-individual variance estimates; variance components are reported on the log-activity scale. Each cell within a heatmap represents a significant estimate ($\alpha < .05$) from a separate model fitted to data from the pair of days indicated by the axes. The horizontal axis gives the first day and the vertical axis gives the second day; only unique day pairs are shown. Cells on the diagonal therefore represent models fitted to observations from a single day, whereas off-diagonal cells represent models fitted to observations from two different days. Cell color denotes the estimated value. At the daily aggregation level, estimates from same-day models cannot be obtained because each individual contributes only one observation per day at this aggregation level. Empty cells elsewhere indicate estimates that did not reach the statistical significance threshold. Repeatability is bounded between 0 and 1. Variance components are not inherently bounded, but all estimated values in this dataset were below 1; therefore, a common color scale is used across panels.

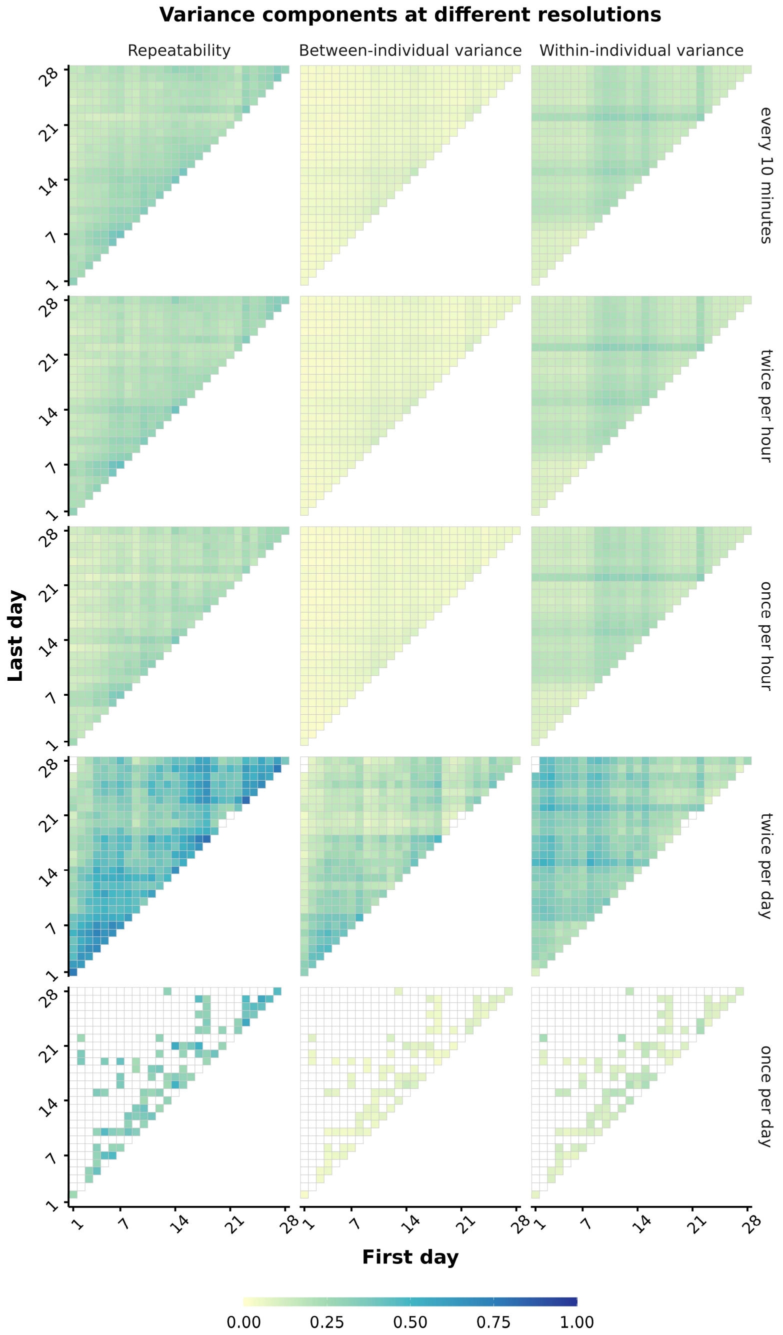


*Figure S20:* Repeatability and variance components across sampling rates computed from activity data tracked at 1 frame per minute. The raw activity observations (recorded at 5 frames per second) were first aggregated across 10-minute observation intervals, resulting in one activity data point per every 10 minutes, per fish. Rows indicate the sampling rate of these 10-minute observation intervals: continuously every 10 minutes, twice per hour, once per hour, twice per day, and once per day. For example, the *twice per hour* sampling rate means that the model included aggregated activity data from two 10-minute observation intervals per hour, per individual. Columns show, from left to right, heatmaps illustrating repeatability ($R$), among-individual variance, and within-individual variance estimates; variance components are reported on the log-activity scale. Each cell within a heatmap represents a significant estimate ($\alpha < .05$) from a separate model fitted to data from the pair of days indicated by the axes. The horizontal axis gives the first day and the vertical axis gives the second day; only unique day pairs are shown. Cells on the diagonal therefore represent models fitted to observations from a single day, whereas off-diagonal cells represent models fitted to observations from two different days. Cell color denotes the estimated value. At the daily aggregation level, estimates from same-day models cannot be obtained because each individual contributes only one observation per day at this aggregation level. Empty cells elsewhere indicate estimates that did not reach the statistical significance threshold. Repeatability is bounded between 0 and 1. Variance components are not inherently bounded, but all estimated values in this dataset were below 1; therefore, a common color scale is used across panels.

<u>Results including body size</u>

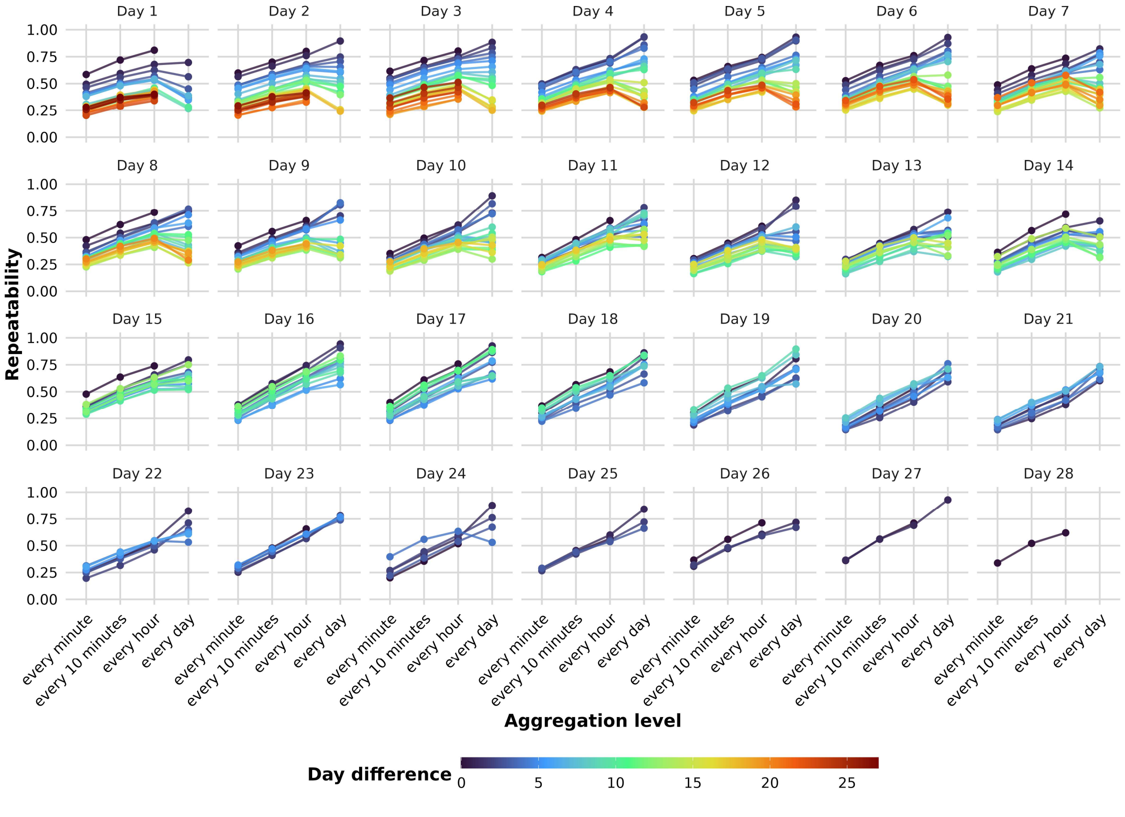

*Figure S21:* Repeatability across aggregation levels computed controlling for body size. Each panel represents a focal day in the fish's life. The vertical axis shows statistically significant ($\alpha < .05$) repeatability estimates. Each filled circle corresponds to one model fitted to data from that focal day and, except where the day difference was zero, some subsequent day (separated from a focal day by the indicated day difference). Colors of circles and connecting lines indicate the number of days separating the two measurement days (day difference; 0–27 days). The horizontal axis indicates the temporal aggregation level applied to the activity data: every minute, every 10 minutes, every hour, or every day. For example, at the *every minute* level, raw activity observations (recorded at 5 frames per second) were averaged within each minute, yielding one observation per individual per minute. A day difference of zero denotes estimates based on data from the focal day alone. Repeatability could not be estimated at the daily aggregation level for these models because this aggregation yields only one observation per individual per day. Other missing estimates indicate that the repeatability estimate did not reach the statistical significance threshold.

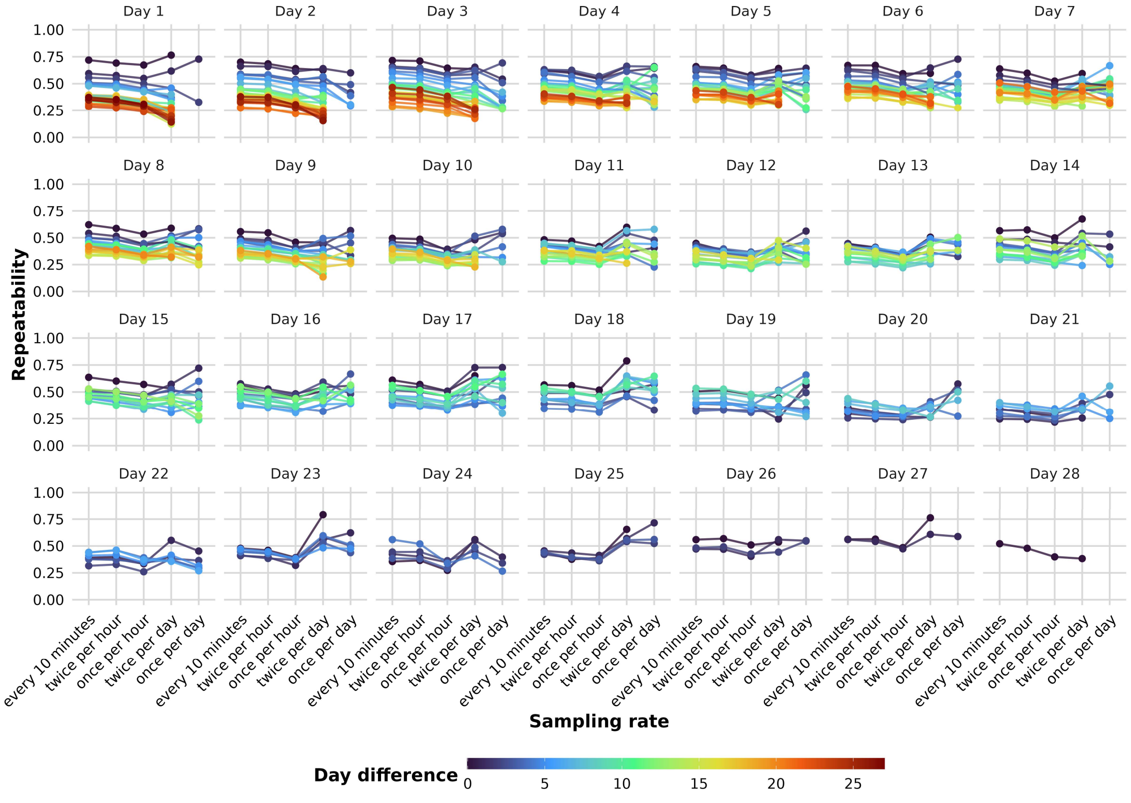


*Figure S22:* Repeatability across sampling rates computed controlling for body size. Each panel represents a focal day in the fish's life. The vertical axis shows statistically significant

($\alpha < .05$) repeatability estimates. Each filled circle corresponds to one model fitted to data from that focal day and, except where the day difference was zero, some subsequent day (separated from a focal day by the indicated day difference). Colors of circles and connecting lines indicate the number of days separating the two measurement days (day difference; 0–27 days). The raw activity observations (recorded at 5 frames per second) were first aggregated across 10-minute observation intervals, resulting in one activity data point per every 10 minutes, per fish. The horizontal axis indicates the sampling rate of these 10-minute observation intervals: continuously every 10 minutes, twice per hour, once per hour, twice per day, and once per day. For example, the *twice per hour* sampling rate means that the model included aggregated activity data from two 10-minute observation intervals per hour, per individual. A day difference of zero denotes estimates based on data from the focal day alone. Repeatability could not be estimated at the once per day sampling rate for these models because this rate yields only one observation per individual, per day. Other missing estimates indicate that the repeatability estimate did not reach the statistical significance threshold.

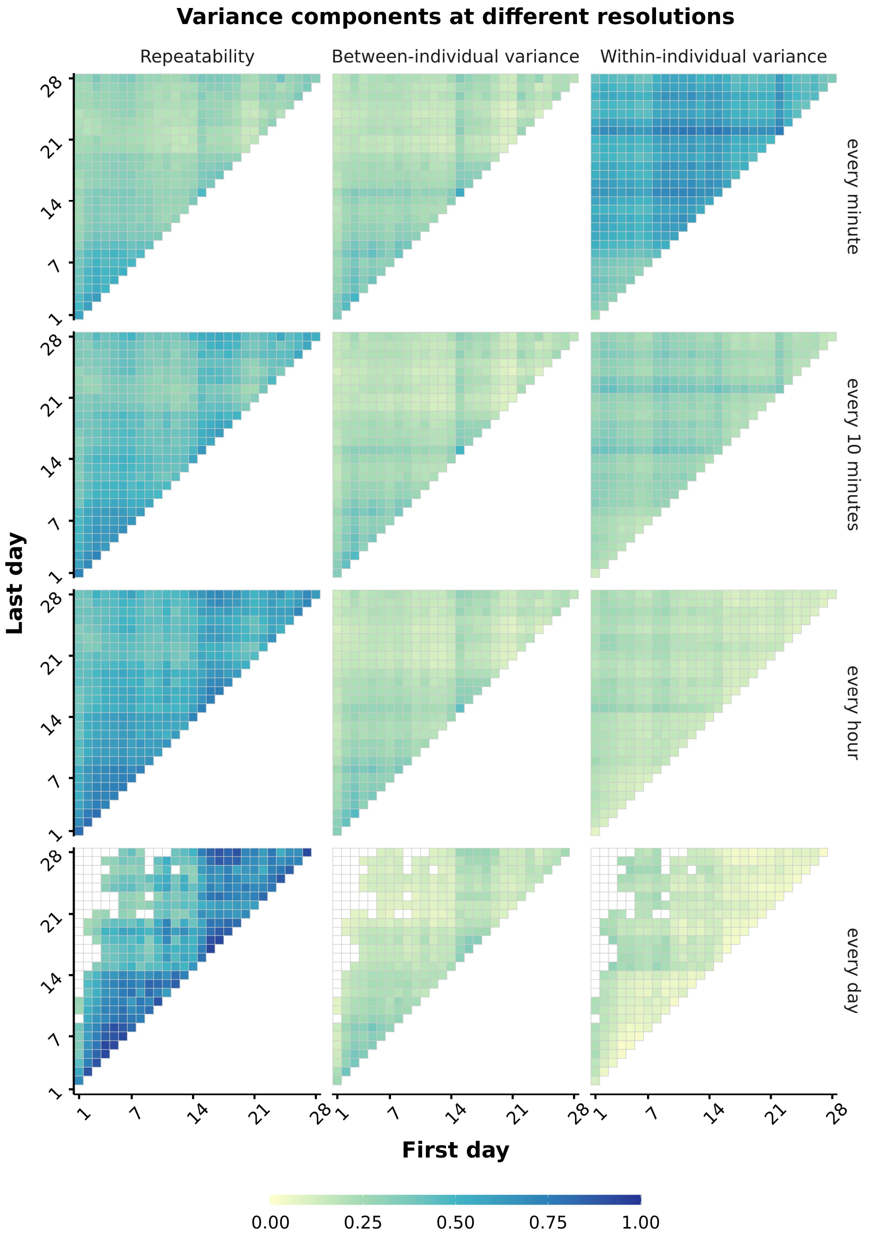


*Figure S23:* Repeatability and variance components across aggregation levels computed controlling for body size. Rows represent the temporal aggregation level applied to the activity data: every minute, every 10 minutes, every hour, and every day. For example, at the every minute level, raw activity observations (recorded at 5 frames per second) were averaged within each minute, yielding one observation per individual per minute. Columns show, from left to right, heatmaps illustrating repeatability ($R$), among-individual variance, and within-individual variance estimates; variance components are reported on the log-activity scale. Each cell within a heatmap represents a significant estimate ($\alpha < .05$) from a separate model fitted to data from the pair of days indicated by the axes. The horizontal axis gives the first day and the vertical axis gives the second day; only unique day pairs are shown. Cells on the diagonal therefore represent models fitted to observations from a single day, whereas off-diagonal cells represent models fitted to observations from two different days. Cell color denotes the estimated value. At the daily aggregation level, estimates from same-day models cannot be obtained because each individual contributes only one observation per day at this aggregation level. Empty cells elsewhere indicate estimates that did not reach the statistical significance threshold. Repeatability is bounded between 0 and 1. Variance components are not inherently bounded, but all estimated values in this dataset were below 1; therefore, a common color scale is used across panels.

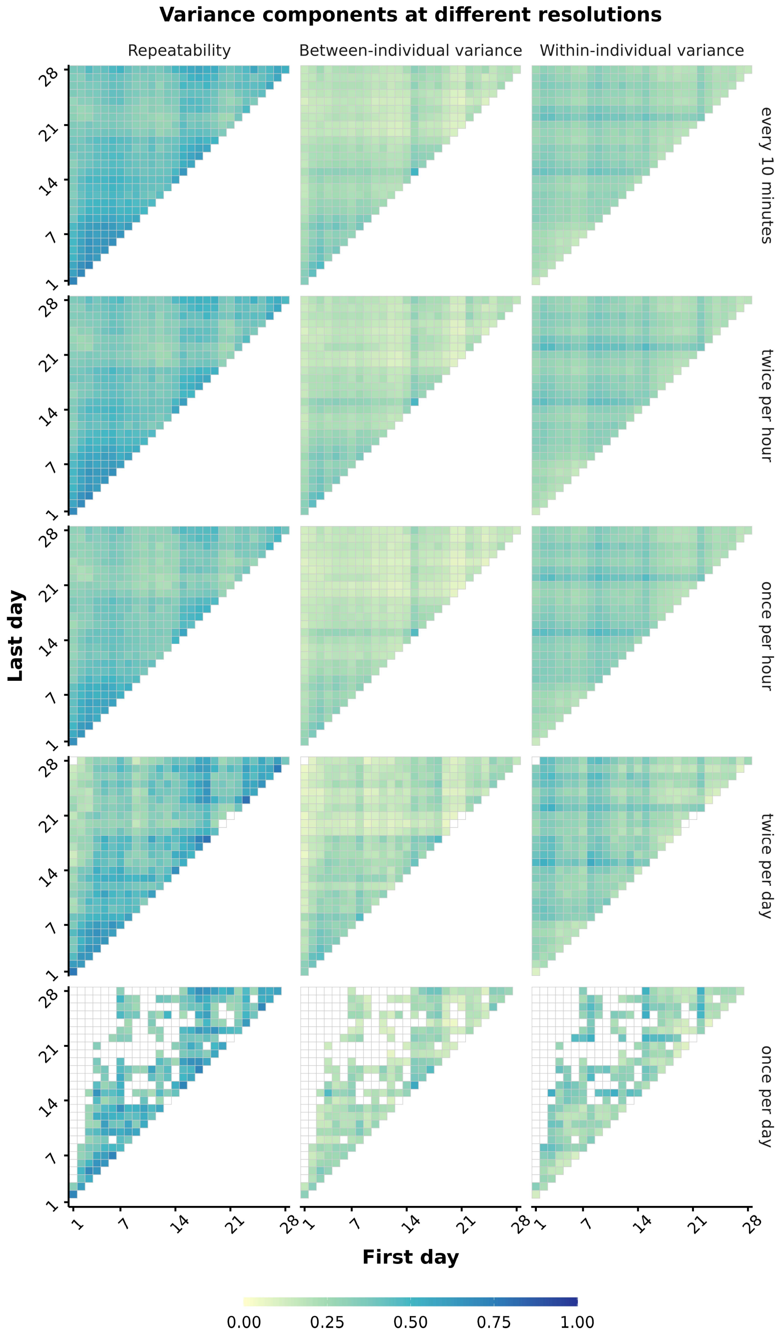


*Figure S24:* Repeatability and variance components across sampling rates. The raw activity observations (recorded at 5 frames per second) were first aggregated across 10-minute observation intervals, resulting in one activity data point per every 10 minutes, per fish. Rows indicate the sampling rate of these 10-minute observation intervals: continuously every 10 minutes, twice per hour, once per hour, twice per day, and once per day. For example, the *twice per hour* sampling rate means that the model included aggregated activity data from two 10-minute observation intervals per hour, per individual. Columns show, from left to right, heatmaps illustrating repeatability ($R$), among-individual variance, and within-individual variance estimates; variance components are reported on the log-activity scale. Each cell within a heatmap represents a significant estimate ($\alpha < .05$) from a separate model fitted to data from the pair of days indicated by the axes. The horizontal axis gives the first day and the vertical axis gives the second day; only unique day pairs are shown. Cells on the diagonal therefore represent models fitted to observations from a single day, whereas off-diagonal cells represent models fitted to observations from two different days. Cell color denotes the estimated value. At the daily aggregation level, estimates from same-day models cannot be obtained because each individual contributes only one observation per day at this aggregation level. Empty cells elsewhere indicate estimates that did not reach the statistical significance threshold. Repeatability is bounded between 0 and 1. Variance components are not inherently bounded, but all estimated values in this dataset were below 1; therefore, a common color scale is used across panels.
